# Regulation of the transcription factor CdnL promotes adaptation to nutrient stress in *Caulobacter*

**DOI:** 10.1101/2023.12.20.572625

**Authors:** Erika L. Smith, Gaël Panis, Selamawit Abi Woldemeskel, Patrick H. Viollier, Peter Chien, Erin D. Goley

## Abstract

In response to nutrient deprivation, bacteria activate a conserved stress response pathway called the stringent response (SR). During SR activation in *Caulobacter crescentus*, SpoT synthesizes the secondary messengers (p)ppGpp, which affect transcription by binding RNA polymerase to downregulate anabolic genes. (p)ppGpp also impacts expression of anabolic genes by controlling the levels and activities of their transcriptional regulators. In *Caulobacter*, a major regulator of anabolic genes is the transcription factor CdnL. If and how CdnL is controlled during the SR and why that might be functionally important is unclear. Here, we show that CdnL is downregulated post-translationally during starvation in a manner dependent on SpoT and the ClpXP protease. Inappropriate stabilization of CdnL during starvation causes misregulation of ribosomal and metabolic genes. Functionally, we demonstrate that the combined action of SR transcriptional regulators and CdnL clearance allows for rapid adaptation to nutrient repletion. Moreover, cells that are unable to clear CdnL during starvation are outcompeted by wild-type cells when subjected to nutrient fluctuations. We hypothesize that clearance of CdnL during the SR, in conjunction with direct binding of (p)ppGpp and DksA to RNAP, is critical for altering the transcriptome in order to permit cell survival during nutrient stress.

**SIGNIFICANCE:** The stringent response (SR) is a ubiquitous bacterial stress response that promotes adaptation to nutrient deprivation. While it is known that SR activation affects RNA polymerase activity to reprogram the transcriptome, the impact of the SR on other transcriptional regulators is not well understood. Here, we show that a conserved transcription factor, CdnL, is cleared upon activation of the SR, and that its clearance is important for cells to efficiently adapt to nutrient fluctuations. Our results suggest that CdnL regulation enables adaptation by transcriptionally downregulating ribosome biosynthesis and flux through metabolic pathways, thereby promoting survival during nutrient stress. As CdnL homologs are broadly found, we hypothesize that CdnL regulation is a conserved mechanism of bacterial adaptation to stress.

## INTRODUCTION

Bacterial replication requires available nutrients and transcription of anabolic genes. When nutrients become limiting, cells adapt to their environment by activating the broadly conserved bacterial stress response pathway known as the stringent response (SR) (1). During the SR, stressors such as nutrient limitation and/or heat shock are sensed by Rel/Spo homolog (RSH) proteins, which synthesize and/or hydrolyze the second messengers guanosine 5’-diphosphate 3’-diphosphate and guanosine 5’-triphosphate 3’-diphosphate (collectively known as (p)ppGpp) (1, 2). (p)ppGpp broadly impacts DNA replication, transcription, translation, and metabolism by directly or indirectly interacting with various proteins to halt growth until conditions improve (1).

*Caulobacter crescentus* (hereafter *Caulobacter*) is a Gram-negative freshwater α-proteobacterium with a dimorphic life cycle that allows it to exist as a nutrient-seeking swarmer cell or as a reproductive stalked cell (1). Nutrient limitation and (p)ppGpp accumulation slow the swarmer-to-stalked transition, likely to allow the swarmer cell more time to seek out nutrients before differentiating into a stalked cell (1–4). This transition is governed by regulatory proteins, whose levels and activities are impacted by (p)ppGpp upon SR activation (3, 5). The availability of nutrients thus regulates the *Caulobacter* cell cycle, promoting anabolic processes and cell cycle progression in nutrient-rich environments, while favoring catabolic processes and nutrient-seeking behavior when nutrients are lacking.

*Caulobacter* inhabits oligotrophic environments and is regularly exposed to fluctuations in nutrient availability; therefore, cells must balance anabolism with SR activation by readily responding to nutrient deprivation (1, 6). In *Caulobacter*, nutrient limitation is sensed by a single bifunctional RSH protein that is known as SpoT, which synthesizes (p)ppGpp during nutrient stress to initiate the SR (1, 5, 6). Ultimately, (p)ppGpp accumulation causes downregulation of genes required for translation, growth, and division, and upregulation of genes important for responding to stress, thus impacting the activities of proteins involved in both anabolism and SR activation (1, 7–9). A major way in which (p)ppGpp exerts these effects on gene regulation is by binding directly to two sites on RNA polymerase (RNAP). These are referred to as “site 1” and “site 2” in *Escherichia coli*, and both sites are conserved in *Caulobacter* (1, 5–11). Site 2 is formed at the interface between RNAP and the transcription factor DksA, and the binding of (p)ppGpp to this site specifically has been implicated in enhancing the influence of DksA on RNAP and leading to inhibition of rRNA promoter activity and decreased anabolism (1, 7, 9, 12).

(p)ppGpp can also cause a downregulation of anabolic gene transcription by controlling the levels and activities of other transcriptional regulators. In *Caulobacter*, a major regulator of anabolic gene transcription and cell cycle progression is the broadly conserved transcription factor CdnL, which stands for “CarD N-terminal like” as it shares homology with the N-terminal domain of the *Myxococcus xanthus* transcription factor, CarD (13–16). Under nutrient-rich conditions, CdnL directly binds to RNAP and promotes transcription of housekeeping genes and growth (13, 17). Similarly, the *Mycobacterium tuberculosis* CdnL homolog CarD was shown to be an activator of rRNA genes whose levels are positively correlated with mycobacterial growth (17–21). We have previously shown that loss of *Caulobacter* CdnL (Δ*cdnL*) results in slowed growth and altered morphology, as well as downregulation of anabolic genes, such as those involved in macromolecule biosynthesis and ribosome biogenesis (14, 16). These data are further supported by metabolomic analyses showing altered metabolite levels in Δ*cdnL* cells (14).

The physiological changes observed in Δ*cdnL* cells mirror many of those of cells undergoing stringent response activation. Indeed, mycobacterial CarD was recently found to be downregulated during stress conditions, though a link to the SR was not examined (22). These putative connections made us consider if CdnL is regulated in a similar manner during the *Caulobacter* SR. Here, we demonstrate that CdnL is regulated post-translationally during the SR, and that this regulation promotes effective adaptation to changes in nutrient availability by altering anabolic gene transcription.

## RESULTS

### CdnL is cleared during the stringent response in a SpoT-dependent manner

Our prior work indicated that many anabolic genes are regulated in a CdnL-dependent manner, apparently echoing transcriptional changes that occur during the SR. This prompted us to review transcriptomic data from our lab and others to explore a potential relationship between CdnL and the SR. In comparing Δ*cdnL* cells to wild-type (WT) starved cells, we find 213 genes downregulated during carbon starvation are also downregulated in Δ*cdnL*, which is a significant enrichment as determined by hypergeometric probability (Fig. 1A) (14, 23). This enrichment is, at least in part, SpoT-dependent, as 130 genes that are downregulated in Δ*cdnL* cells are downregulated during carbon starvation in a SpoT-dependent manner, which is also statistically significant (Fig. 1A) (6, 14). These overlaps suggest a link between activation of the SR by SpoT and the absence of CdnL.

**Figure 1:**
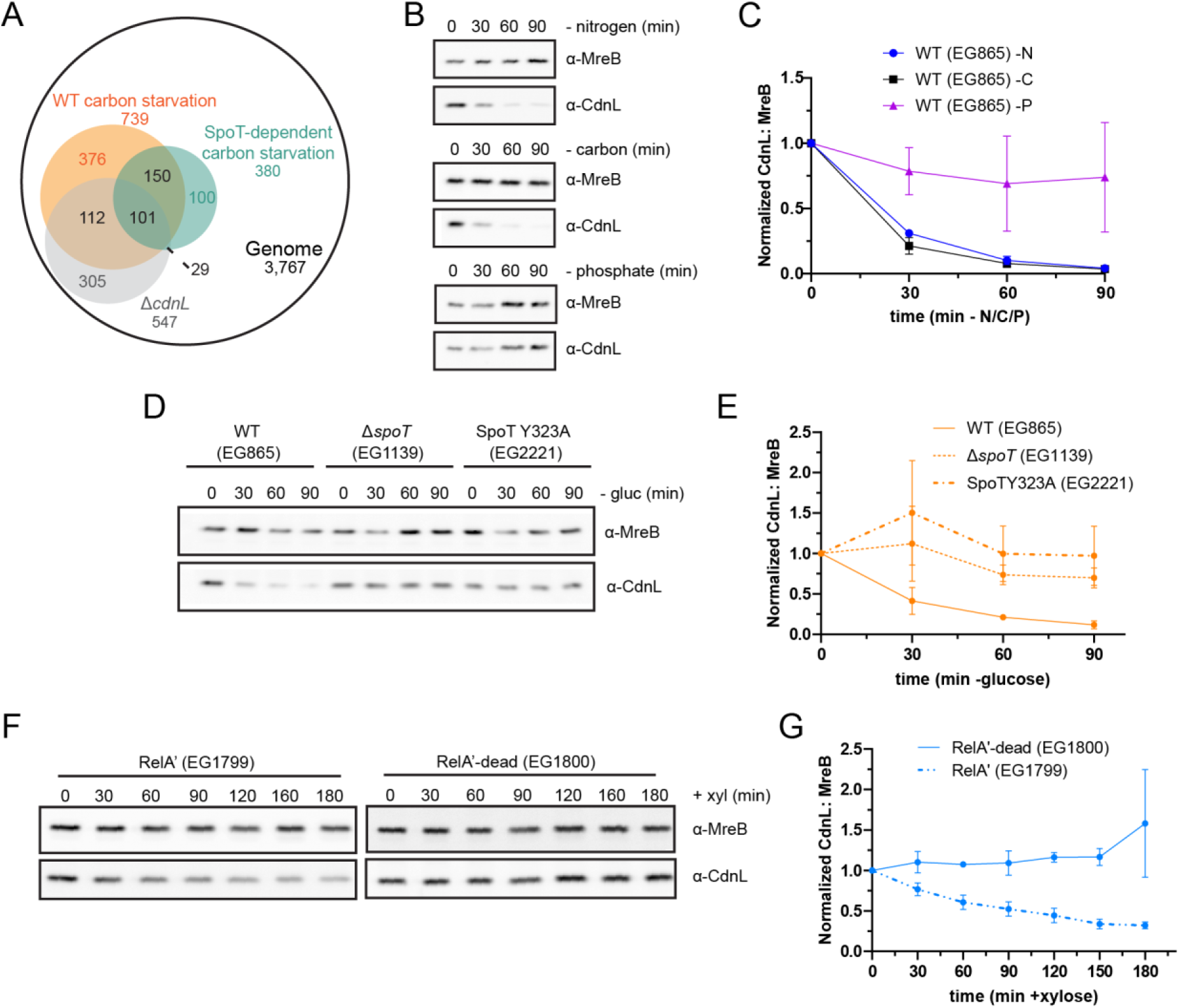
CdnL is cleared during conditions that activate the stringent response. A. Venn diagram of genes downregulated in Δ*cdnL*, in WT carbon starved cells, and in a SpoT-dependent manner (6, 14, 23). 213 genes overlapped between the Δ*cdnL* dataset and the WT carbon starved dataset (14, 23). 130 genes overlapped between the Δ*cdnL* dataset and the SpoT-dependent carbon starved dataset (6, 14). Both are significant enrichments (p < 0.05) as determined by hypergeometric probability. B. Representative western blots of CdnL during 90 minutes of nitrogen, carbon, and phosphate starvation. Protein samples were taken every 30 minutes. MreB was used as a loading control. C. Densitometry of CdnL levels (normalized to MreB) relative to t = 0 during nitrogen (-N), carbon (-C), and phosphate (-P) starvations from western blots as performed in B. Error bars represent +/- 1 SD of 3 biological replicates. D. Representative western blot of CdnL during 90 minutes of glucose (gluc) starvation in a WT, Δ*spoT,* or SpoTY323A background. Protein samples were taken every 30 minutes. MreB was used as a loading control. E. Densitometry of CdnL levels (normalized to MreB) relative to t = 0 from western blots as performed in D. Error bars represent +/- 1 SD of 3 biological replicates. F. Representative western blot of CdnL during 180 minutes of xylose (+xyl) induction of RelA’ or RelA’-dead. Protein samples were taken every 30 minutes. MreB was used as a loading control. G. Densitometry of CdnL levels (normalized to MreB) relative to t = 0 from western blots as performed in F. Error bars represent +/- 1 SD of 3 biological replicates.

To investigate a potential connection between SR activation and loss of CdnL, we assessed CdnL protein levels during nitrogen, carbon, and phosphate starvation. Both carbon and nitrogen limitation activate the SR in *Caulobacter*, while phosphate limitation does not (1). To do this, *Caulobacter* cells were grown in minimal media, then washed and incubated in media lacking a carbon, nitrogen, or phosphate source. Cell lysates were sampled at up to 90 minutes of starvation and CdnL levels were assessed by immunoblotting. We found that CdnL was cleared in WT cells under both nitrogen and carbon starvation with half-lives of 18 and 15 min, respectively (Fig. 1B,C and Table S1). Notably, CdnL levels remained stable under phosphate starvation (Fig. 1B,C and Table S1).

The fact that CdnL is cleared upon carbon and nitrogen starvation, but not phosphate starvation, is consistent with CdnL regulation being downstream of SR activation. Therefore, we asked if CdnL clearance depended on the only RSH enzyme in *Caulobacter*, SpoT, by repeating these starvation experiments in cells lacking SpoT (Δ*spoT)* (6). During nitrogen and carbon starvation, CdnL levels were significantly more stable in the Δ*spoT* background than in WT, with the half-lives increasing to 60 min and over 500 min, respectively (Fig. 1D,E, Fig. S1A,B, and Table S1). CdnL was also stabilized during carbon starvation in the presence of a (p)ppGpp synthetase-dead SpoT variant (SpoTY323A) (Fig. 1D,E and Table S1) (2). These observations indicate that both SpoT and (p)ppGpp are necessary for CdnL clearance during starvation.

We next tested if (p)ppGpp is sufficient for CdnL clearance using a strain harboring a xylose-inducible, constitutively active version of the *E. coli* (p)ppGpp synthetase RelA (RelA’) or a catalytically inactive RelA variant (RelA’-dead) as a negative control (4). We found that CdnL was cleared under nutrient-replete conditions upon expression of RelA’ with a half-life of 103 min, while CdnL is stable in the presence of RelA’-dead, indicating that (p)ppGpp is sufficient to clear CdnL (Fig. 1F,G and Table S1). Taken together, these results suggest that CdnL is cleared during conditions that activate the SR, and that (p)ppGpp is both necessary and sufficient to induce clearance. For simplicity, we chose to use carbon starvation to probe the mechanism and importance of CdnL clearance during stringent activation.

### Transcriptional control of *cdnL* is not sufficient to regulate CdnL levels during the SR

We observed that CdnL protein levels decrease upon carbon starvation, but it was unclear if regulation of CdnL levels during starvation occurs transcriptionally, post-transcriptionally, and/or post-translationally. Indeed, previous studies showed that *cdnL* transcript levels also decrease upon carbon starvation (23). We first asked if transcriptional activity at the *cdnL* promoter contributes significantly to controlling CdnL levels during the SR. To do this, we expressed *cdnL* from a xylose-inducible promoter in Δ*cdnL*, Δ*spoT*Δ*cdnL*, and SpoTY323A Δ*cdnL* backgrounds. In this system, we can shut off *cdnL* transcription independently of SR activation or (p)ppGpp production by washing out xylose and resuspending cells in nutrient-replete media lacking xylose (M2G), or in carbon starvation media lacking both xylose and glucose (M2). Under carbon starvation conditions and after the removal of xylose to prevent further *cdnL* transcription, CdnL was rapidly cleared in Δ*cdnL* cells with a half-life of 13 min. In contrast, for Δ*spoT*Δ*cdnL* and SpoTY323A Δ*cdnL* cells, CdnL levels remained stable with half-lives of 139 and 334 min, respectively (Fig. S2A,B and Table S1). This suggests that transcriptional control is not sufficient for CdnL clearance. There appear to be additional factors governing CdnL stability during the SR, however. While CdnL was cleared more rapidly in starvation conditions compared to nutrient-replete conditions in a Δ*cdnL* background after the removal of xylose (13 min compared to 27 min), we found that in the Δ*spoT*Δ*cdnL* and SpoTY323A Δ*cdnL* backgrounds, CdnL was actually cleared more rapidly in nutrient-replete conditions compared to starvation conditions after the removal of xylose (70 and 75 min compared to 139 and 334 min, respectively) (Fig. S2A-D and Table S1). These data imply that CdnL levels are regulated post-transcriptionally in a manner that is partially SpoT-dependent.

### CdnL is a ClpXP target

Since transcriptional regulation is not sufficient to control CdnL levels during the SR, we next asked if CdnL is regulated post-translationally. CdnL bears two alanine residues on the C-terminus, which is a common degradation signal (degron) recognized by the ClpXP protease (24, 25). It was previously shown that the CdnL C-terminus was important for ClpXP-mediated degradation of CdnL *in vivo*, which caused us to wonder how this might relate to regulation of CdnL during starvation (13). We asked if eliminating this putative degron impacts CdnL levels during starvation by changing the two alanine residues to aspartate residues, thus creating the variant CdnLDD (25). We found that CdnLDD protein was not cleared during either nitrogen or carbon starvation, suggesting that ClpXP targets CdnL for degradation during the SR (Fig. 2A,B and Fig. S1A,C). We asked if CdnL is a direct proteolytic target of ClpXP *in vitro* and found that purified ClpXP was capable of degrading purified CdnL, but not purified CdnLDD (Fig. 2C). The ability of ClpXP to degrade CdnL *in vitro* is not affected by (p)ppGpp, as the addition of (p)ppGpp did not further stimulate CdnL turnover. Finally, we assessed if ClpX activity was required *in vivo* for CdnL clearance during carbon starvation using a dominant-negative ATPase-dead ClpX variant (ClpX*) (26, 27). When cells were starved of carbon after a 1-hour induction of ClpX*, CdnL levels were once again stabilized (Fig. 2D,E). These data confirm that CdnL is a ClpXP target as well as demonstrate that ClpXP-mediated proteolysis is required for regulation of CdnL during carbon starvation.

**Figure 2:**
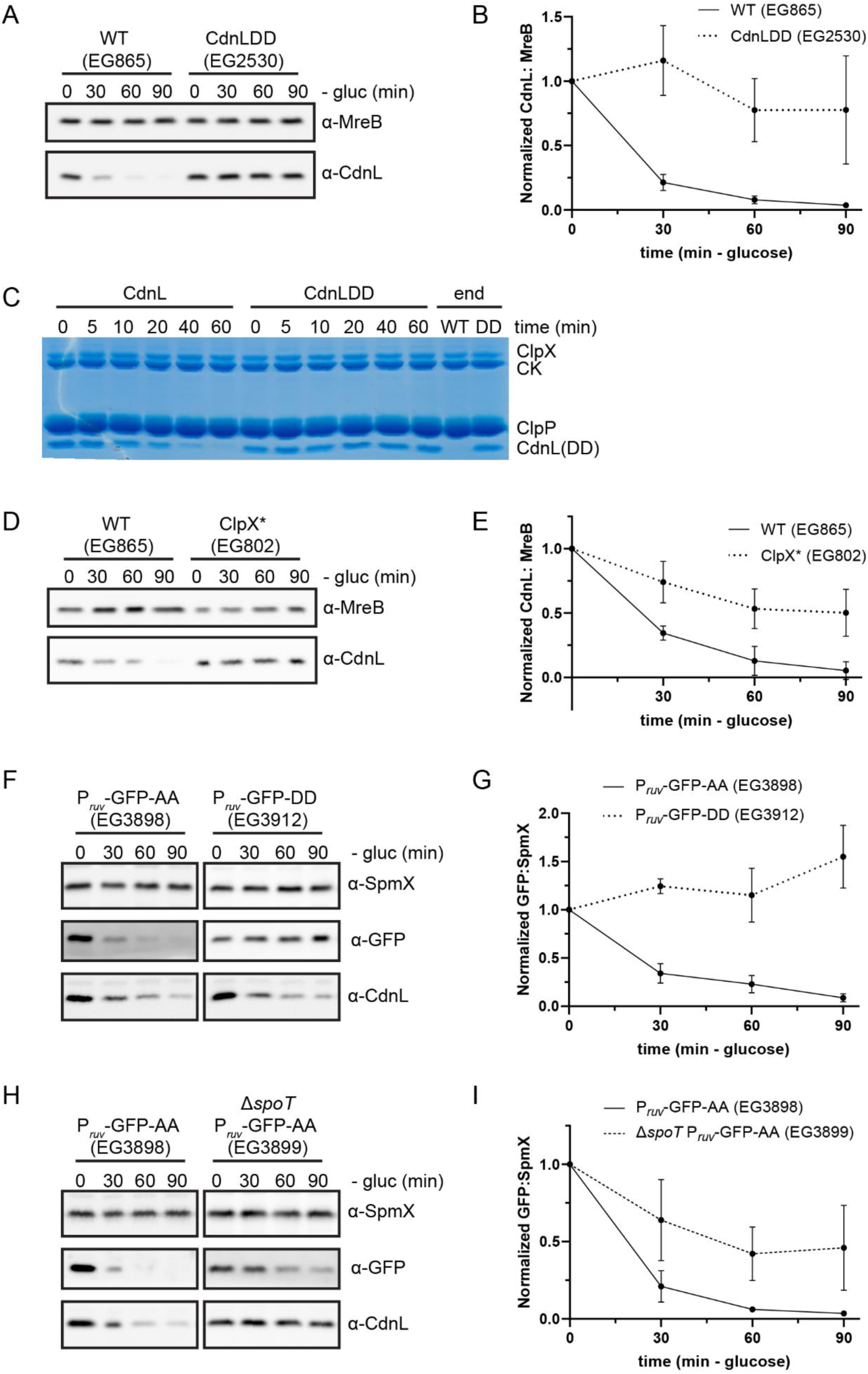
ClpXP is implicated in CdnL clearance. A. Representative western blot of CdnL or CdnLDD during 90 minutes of glucose (gluc) starvation. Protein samples were taken every 30 minutes. MreB was used as a loading control. B. Densitometry of CdnL or CdnLDD levels (normalized to MreB) relative to t = 0 from western blots as performed in B. Error bars represent +/- 1 SD of 3 biological replicates. C. *In vitro* degradation of CdnL or CdnLDD. Protein samples were taken at indicated time points. CK indicated creatine kinase, which is used for ATP regeneration in the reaction. D. Representative western blot of CdnL during 90 minutes of glucose (gluc) starvation following a 2-hour induction of ClpX* (EG802) with 0.3% xylose. Protein samples were taken every 30 minutes. MreB was used as a loading control. E. Densitometry of CdnL levels (normalized to MreB) relative to t = 0 from western blots as performed in D. Error bars represent +/- 1 SD of 3 biological replicates. F. Representative western blot of GFP-AA or GFP-DD during 90 minutes of glucose (gluc) starvation. Protein samples were taken every 30 minutes. SpmX was used as a loading control. CdnL was used as a comparison. G. Densitometry of GFP-AA or GFP-DD levels (normalized to SpmX) relative to t = 0 from western blots as performed in F. Error bars represent +/- 1 SD of 3 biological replicates. H. Representative western blot of GFP-AA in a WT or Δ*spoT* background during 90 minutes of glucose (gluc) starvation. Protein samples were taken every 30 minutes. SpmX was used as a loading control. CdnL was used as a comparison. I. Densitometry of GFP-AA levels (normalized to SpmX) relative to t = 0 from western blots as performed in H. Error bars represent +/- 1 SD of 3 biological replicates.

It is evident that the CdnL C-terminus is critical for the regulation of CdnL by ClpXP. To assess if the CdnL C-terminus is sufficient for ClpXP-mediated degradation during starvation, we tagged GFP with the last 15 amino acids from the CdnL tail (GFP-AA) or with the last 15 amino acids from the stabilized CdnL tail (GFP-DD). We then expressed these constructs on a plasmid under control of the *ruv* operon promoter, as transcript levels for the *ruvA* DNA helicase gene are not affected during carbon starvation (6). Strikingly, GFP-AA was rapidly degraded during carbon starvation while GFP-DD remained stable, demonstrating that the CdnL tail is indeed sufficient for degradation of GFP (Fig. 2F,G). We next sought to test if there is a SpoT-dependence to this degradation. To do this, we put the GFP-AA construct into the Δ*spoT* background and repeated the carbon starvation. GFP-AA was degraded more slowly in a Δ*spoT* background than in WT, with a half-life of 68 min in comparison to the 14 min half-life in the WT background, indicating that there is at least a partial SpoT-dependence to ClpXP-mediated degradation (Fig. 2H,I and Table S1).

### The CdnL-RNAP interaction is not sufficient to protect CdnL from proteolysis

There appear to be other factors governing CdnL stability outside of the C-terminus since we found that GFP-AA was cleared, albeit slowly, in Δ*spoT* cells, while CdnL itself remained stable (Fig. 2H,I and Fig. 1D,E). Since CdnL interacts with RNAP while GFP does not, we hypothesized that the interaction with RNAP may reduce CdnL degradation, possibly by protecting CdnL from proteolysis by ClpXP. To assess this, we constructed two CdnL point mutants, V39A and P54A, that have a reduced affinity for RNAP (13). We then assessed their stabilities during carbon starvation. Both the V39A and P54A mutants exhibited reduced half-lives (15 and 18 min, respectively) compared to WT CdnL (32 min) (Fig. S3A,B and Table S1). These observations are consistent with the interaction between CdnL and RNAP contributing to CdnL stability under carbon starvation conditions. To test the sufficiency of this interaction in determining CdnL stability, we put the V39A and P54A point mutants into a Δ*spoT* background and performed another carbon starvation. If the CdnL-RNAP interaction serves as a barrier to CdnL proteolysis by ClpXP, we expected that these point mutants with a reduced RNAP interaction may still be less stable than WT CdnL in a Δ*spoT* background. However, we found that the point mutants are equally as stable as WT CdnL in the Δ*spoT* background (Fig. S3C,D). These results indicate that disrupting the RNAP-CdnL interaction is not sufficient to promote CdnL clearance during starvation and reiterates the dependency on SpoT to stimulate CdnL turnover.

### CdnL is cleared during stationary phase in a SpoT and ClpXP independent manner

In addition to starvation, another time in which cells experience nutrient stress is during stationary phase when resources become depleted as a population saturates. (p)ppGpp levels peak upon entry into stationary phase, and then dip to stabilize at levels that are still higher than basal levels occurring during logarithmic growth (28). As we have found that both SpoT and ClpXP play important roles in maintaining CdnL levels during starvation when (p)ppGpp levels are high, we were curious to understand how these factors might contribute to regulating CdnL levels during stationary phase. To this end, we assessed CdnL protein levels from mid-log to late stationary phase in WT, CdnLDD, and Δ*spoT* (Fig. S4). CdnLDD levels were consistently higher than CdnL levels in both WT and Δ*spoT*, with Δ*spoT* having the lowest CdnL levels throughout. In all strains, CdnL levels started to decline at an OD_600_ of about 1.0. This reduction was stark for both WT and Δ*spoT*, while it was more gradual for CdnLDD. Surprisingly, we found that CdnL was cleared in all strains by an OD_600_ of about 1.6, suggesting this mechanism of regulation is (p)ppGpp-independent and distinct from that of carbon starvation.

### CdnL and CdnLDD exhibit reduced chromosomal binding during starvation

Mechanistically, we have shown that CdnL is cleared post-transcriptionally in a SpoT- and ClpXP-dependent manner, and that mutation of the C-terminal degradation signal to CdnLDD stabilizes protein levels during starvation. As CdnL is a transcription factor, we sought to understand if stabilization of CdnL protein would permit increased CdnL occupancy on the chromosome during starvation, presumably via interaction with RNAP, thus leading to transcriptomic changes that would be disadvantageous during carbon starvation. To begin to understand the consequences of CdnL regulation on transcriptional reprogramming during nutrient limitation, we used chromatin immunoprecipitation sequencing (ChIP-seq) to assess the occupancy of WT CdnL and CdnLDD on the chromosome under both nutrient-replete (M2G) conditions and after 60 minutes of carbon starvation. We selected this time point as WT CdnL was almost completely cleared (Fig. 1B,C) and broad transcriptional changes are reported at this point (23). After identifying 107 peaks with a > 2-fold enrichment in the binding profile of WT CdnL in M2G compared to the Δ*cdnL* control, we looked at these peaks in the binding profiles of WT CdnL and CdnLDD in both M2G and M2 (Fig. S5 and Dataset S1). In M2G, the binding profiles of WT CdnL and CdnLDD were well correlated with each other and with our prior ChIP-Seq analysis of CdnL association across the genome (Fig. 3A) (14). Consistent with our previous transcriptomic analyses, DAVID analysis of locus function revealed a significant enrichment in aminoacyl tRNA biosynthesis and ribosomal genes (Dataset S1) (14). During carbon starvation (M2 media), we observed three clusters of loci with distinct CdnL association profiles. WT CdnL had an overall reduction in binding at Group 1 genomic loci, while Groups 2 and 3 showed either equal or increased CdnL binding (Fig. 3A). This is surprising given that CdnL protein levels are significantly reduced in the absence of carbon (Fig. 1B,C). Interestingly, CdnLDD showed an overall decrease in binding after 60 minutes of starvation, similar to WT, despite the fact that CdnLDD levels in M2 are comparable to WT CdnL levels in M2G (Fig. 3A). These observations suggest that while CdnLDD is highly stable, it largely does not associate with the DNA after 60 minutes of carbon starvation.

**Figure 3:**
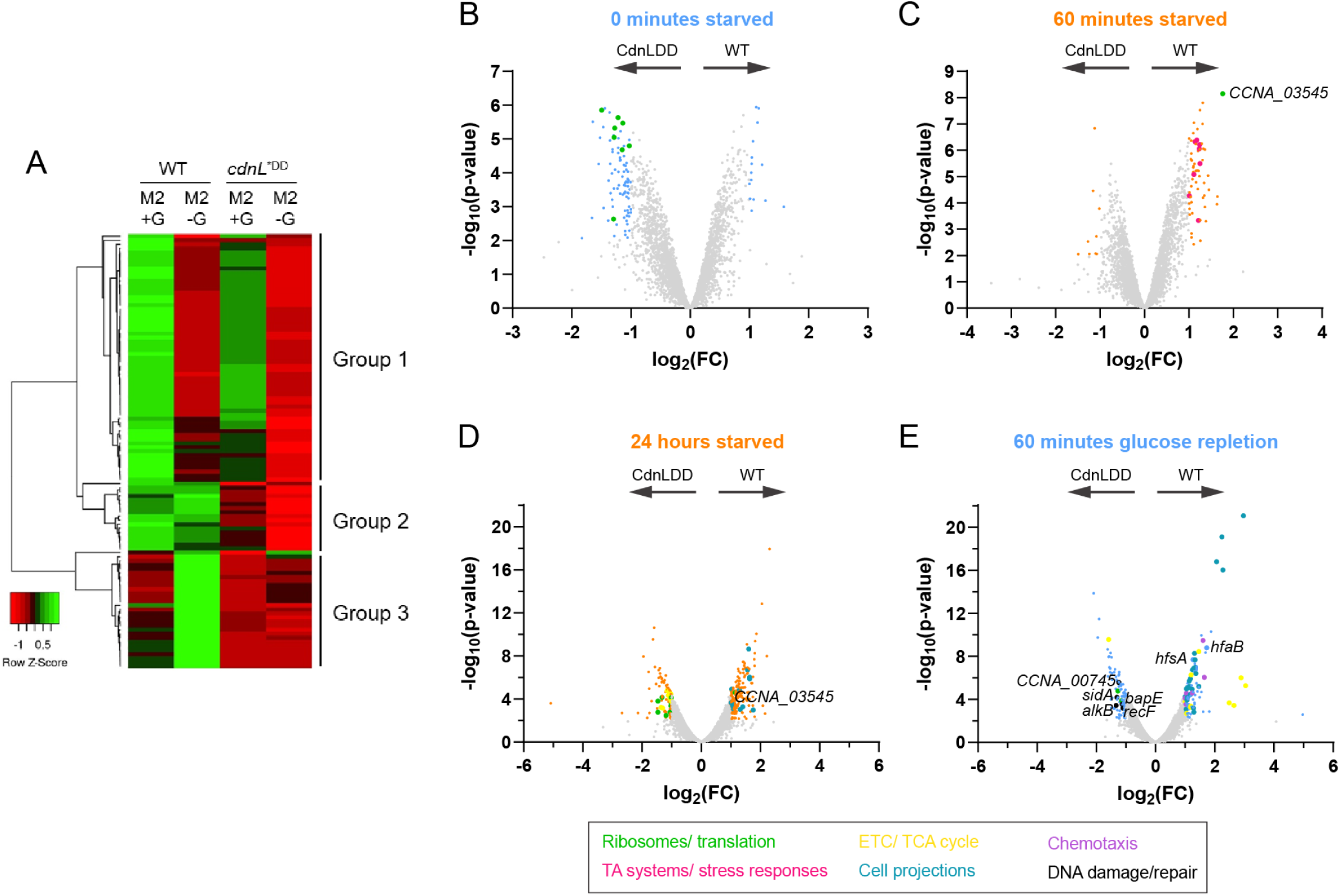
CdnL stabilization impacts transcription of ribosome-related genes. A. Heat map of CdnL/CdnLDD binding across the chromosome in M2G and 60 minutes of glucose starvation in M2. Green indicates high relative binding, while red indicates low relative binding. Group 1 shows genomic loci with a reduction of WT CdnL binding after starvation in M2. Group 2 shows genomic loci with equal WT CdnL binding after starvation in M2. Group 2 shows genomic loci with an increase in WT CdnL binding after starvation in M2. Row Z-score = (CdnL Fold enrichment value in the sample of interest - Mean CdnL Fold enrichment across all samples) / Standard Deviation. B. Volcano plot comparing differences in gene expression between WT and CdnLDD at 0 minutes of starvation, as measured by RNAseq. Negative log_10_ of the p-value is plotted against log_2_ of the fold change mRNA counts in WT vs CdnLDD (FC = WT mRNA counts/ CdnLDD mRNA counts). Light blue points indicate transcripts with FDR < 0.05 and |log_2_(FC)| > 1, while gray points indicate transcripts that are not significantly different. Green points indicate transcripts associated with ribosomes and translation. Arrows indicate direction of higher expression for respective strains. C. Volcano plot comparing differences in gene expression between WT and CdnLDD at 60 minutes of starvation, as measured by RNAseq. Negative log_10_ of the p-value is plotted against log_2_ of the fold change mRNA counts in WT vs CdnLDD (FC = WT mRNA counts/ CdnLDD mRNA counts). Orange points indicate transcripts with FDR < 0.05 and |log_2_(FC)| > 1, while gray points indicate transcripts that are not significantly different. Pink points indicate transcripts associated with toxin-antitoxin (TA) systems and stress responses. Arrows indicate direction of higher expression for respective strains. D. Volcano plot comparing differences in gene expression between WT and CdnLDD at 24 hours of starvation, as measured by RNAseq. Negative log_10_ of the p-value is plotted against log_2_ of the fold change mRNA counts in WT vs CdnLDD (FC = WT mRNA counts/ CdnLDD mRNA counts). Orange points indicate transcripts with FDR < 0.05 and |log_2_(FC)| > 1, while gray points indicate transcripts that are not significantly different. Teal points indicate transcripts associated with cell projections, such as pili and flagella, while yellow points indicate transcripts associated with the electron transport chain (ETC) and tricarboxylic acid (TCA) cycle. Arrows indicate direction of higher expression for respective strains. E. Volcano plot comparing differences in gene expression between WT and CdnLDD at 0 minutes of starvation, as measured by RNAseq. Negative log_10_ of the p-value is plotted against log_2_ of the fold change mRNA counts in WT vs CdnLDD (FC = WT mRNA counts/ CdnLDD mRNA counts). Light blue points indicate transcripts with FDR < 0.05 and |log_2_(FC)| > 1, while gray points indicate transcripts that are not significantly different. Purple points indicate transcripts associated with chemotaxis, while black points indicate transcripts associated with DNA damage and repair. Arrows indicate direction of higher expression for respective strains.

### CdnL stabilization alters ribosomal and metabolic gene transcription

Although CdnLDD does not interact with the chromosome after 60 minutes of carbon starvation, we were interested in understanding if CdnLDD could still facilitate transcriptional changes. To address this, we used RNA-sequencing (RNA-seq) to assess the transcriptional profiles of WT and CdnLDD at 0 and 60 minutes of carbon starvation. In comparing CdnLDD to WT at 0 minutes carbon starvation, we found that 101 genes were at least 2-fold differentially regulated (Fig. 3B and Dataset S2). This number is smaller than the 525 genes that we had previously found to be differentially regulated when comparing WT to Δ*cdnL*; however, Δ*cdnL* cells have growth and morphological defects under nutrient-rich conditions, while CdnLDD cells do not exhibit an obvious phenotype (14). DAVID functional annotation analyses pointed to an enrichment of transcripts related to the ribosome in CdnLDD, which reinforces our ChIP-seq data and previous transcriptomic analyses implicating CdnL in the regulation of anabolic genes (Fig. 3A and Dataset S2) (14).

At 60 minutes of starvation, 71 genes were found to be at least 2-fold differentially regulated between WT and CdnLDD (Fig. 3C and Dataset S3). This set of genes showed no overlap with the 101 genes differentially regulated at 0 minutes of starvation. The most highly upregulated transcript in WT compared to CdnLDD was CCNA_03545 (ribosome silencing factor), which acts to separate the large and small ribosomal subunits during stationary phase and stress, further implicating CdnLDD in altered ribosome activity (Fig. 3C and Dataset S3) (29). Additionally, DAVID functional annotation analyses indicated a relative upregulation of transcripts associated with toxin-antitoxin systems and stress responses in WT, suggesting that CdnLDD may not be appropriately responding to nutrient deprivation (Fig. 3C and Dataset S3). Nonetheless, the relatively small differences in the transcriptional profiles between WT and CdnLDD is consistent with their similar binding profiles demonstrated by ChIP-seq and further suggests that CdnL stabilization does not cause global changes in transcription after 60 minutes of starvation.

While CdnL stabilization does not globally impact the immediate transcriptional response to starvation, we wondered how the transcriptome might be impacted over a longer starvation. It is possible that *Caulobacter* would experience even longer periods of starvation in nature. Indeed, we found that the WT transcriptome changes from 60 minutes of starvation to 24 hours of starvation, prompting us to investigate potential differences between WT and CdnLDD transcripts at additional time points (Dataset S4). We decided to assess the WT and CdnLDD transcriptional profiles at 24 hours of starvation, a time point in which CdnLDD remains stable (Fig. S6A). At this time point, we found 279 genes to be 2-fold differentially regulated between WT and CdnLDD, 21 of which were also differentially expressed after 60 minutes (Fig. 3D and Dataset S5). Of these genes, 177 were more highly expressed in WT, including the ribosome silencing factor again, while 102 were more highly expressed in CdnLDD (Fig. 3D and Dataset S5). For the genes more highly expressed in WT, DAVID functional annotation analyses revealed a relative upregulation of transcripts associated with cell projections, such as flagella and pili (Fig. 3D and Dataset S5). Lower relative expression of these transcripts in CdnLDD could suggest that CdnLDD is continuing to promote cell cycle progression in spite of (p)ppGpp accumulation, which normally slows the swarmer-to-stalked transition and leads to an accumulation of flagellated swarmer cells (2). Of the genes upregulated in CdnLDD compared to WT, many were associated with ribosomes, including translation initiation and elongation factors (Fig. 3D and Dataset S5). The production of ribosomes, and protein synthesis in general, is typically reduced under starvation conditions to slow growth and allow resources to be diverted to amino acid biosynthesis (30–32). Additionally, transcripts associated with metabolic pathways, such as the electron transport chain (ETC) and tricarboxylic acid (TCA) cycle, were found to be enriched in CdnLDD, suggesting increased flux through these pathways (Fig. 3D and Dataset S5).

We also wanted to understand if CdnL stabilization could affect transcription during the adaptation phase when nutrients are replenished after a long starvation. Therefore, we assessed the transcriptome of WT and CdnLDD 60 minutes after the addition of glucose, a time point in which WT CdnL has not returned to basal levels (Fig. S6A). After replenishing glucose and allowing the cells to recover for 60 minutes, 199 genes were found to be differentially regulated (Fig. 3E and Dataset S6). In addition to again finding a relative upregulation of transcripts associated with cell projections in WT, DAVID analyses also showed an enrichment of transcripts for genes involved in chemotaxis and oxidative phosphorylation (Fig. 3E and Dataset S6). Transcripts associated with holdfast synthesis (*hfsA*) and attachment (*hfsB*) were also upregulated (Fig. 3E and Dataset S6). Together, these observations suggest that the WT cells were responding to nutrient availability by resuming growth, cell cycle progression, and polar development. For CdnLDD, ribosome-related transcripts were again found to be upregulated (Fig. 3E and Dataset S6). Interestingly, additional transcripts enriched in CdnLDD point to an upregulation of genes involved in the DNA damage response and repair pathways, including the alkylated DNA repair protein *alkB* (CCNA_00009), the SOS-induced inhibitor of cell division *sidA* (CCNA_02004), an oxidative DNA demethylation family protein (CCNA_00745), the bacterial apoptosis endonuclease *bapE* (CCNA_00663), and the DNA replication and repair protein *recF* (CCNA_00158) (Fig. 3E and Dataset S6). Upregulation of these transcripts could suggest DNA damage in CdnLDD.

In summary, these RNA-seq analyses reveal that CdnL stabilization causes increased levels of transcripts associated with ribosomal and protein synthesis genes under both nutrient-rich and carbon-starved conditions, as well as an increase in transcripts associated with metabolic pathways after 24 hours of starvation. During nutrient-repletion, CdnL stabilization leads to an upregulation of transcripts related to DNA damage response and repair proteins.

### CdnL clearance during carbon starvation is necessary for efficient outgrowth upon nutrient repletion

Since we observed transcriptional changes between WT and CdnLDD during starvation and upon nutrient repletion, we wondered if CdnL stabilization impacts the ability of cells to adapt to nutrient fluctuations. Being a global regulator of growth, we hypothesized that CdnL clearance during starvation facilitates transcriptional changes that enables adaptation by ultimately promoting survival instead of proliferation. To this end, we measured the growth of WT and CdnLDD strains both before a 24-hour carbon starvation and after the 24-hour starvation upon glucose repletion. Before starvation, WT and CdnLDD showed similar growth rates, with doubling times of 2.1 and 2.2 hours, respectively (Fig. 4A and Table S2). During the outgrowth from 24 hours of starvation, the strains again had similar growth rates; however, CdnLDD had a slightly longer lag time compared to WT and reached an OD of 0.1 after about 9.3 hours as opposed to 8.3 hours, respectively (Fig. 4B and Table S2).

**Figure 4:**
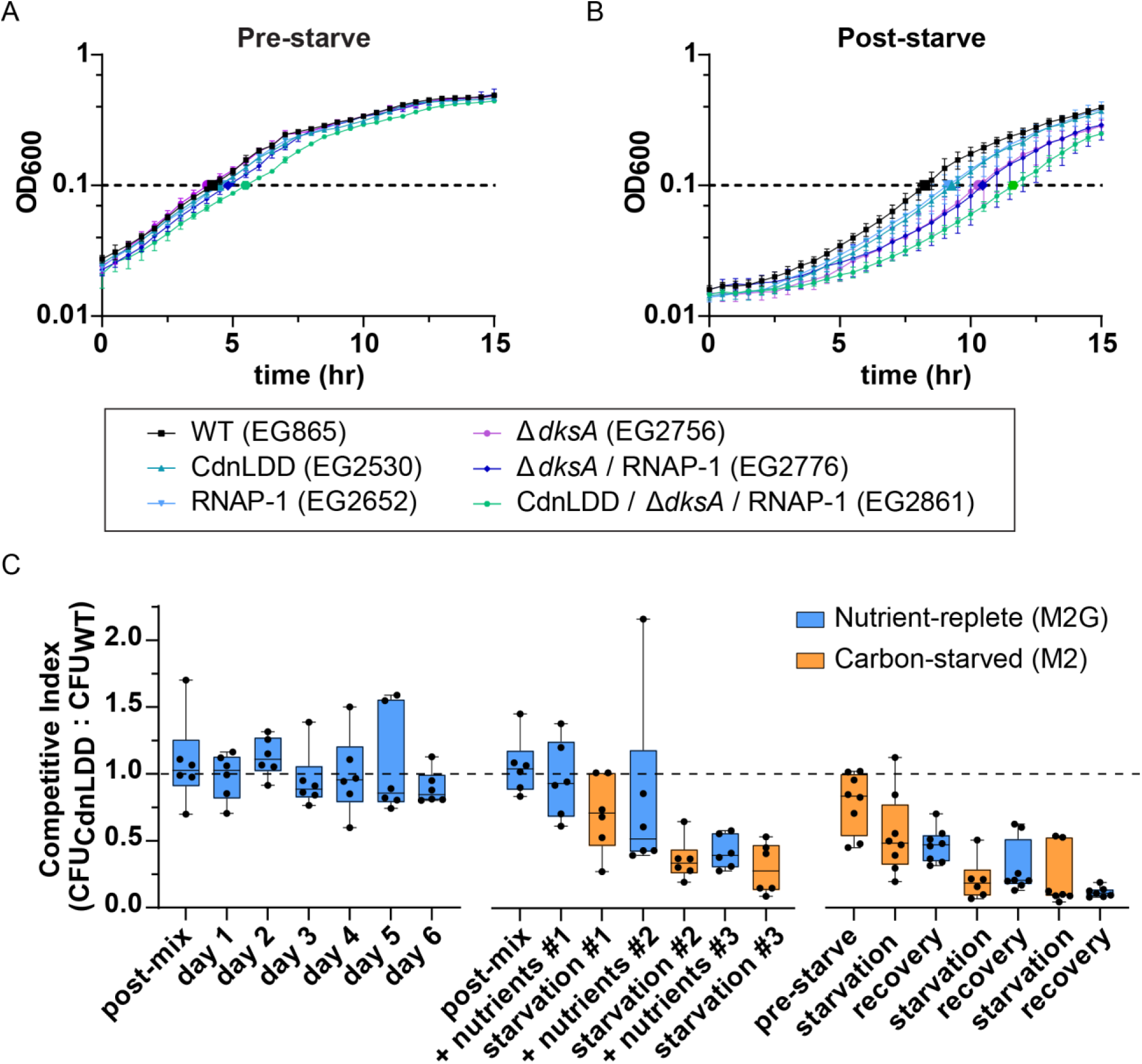
Efficient outgrowth after glucose limitation requires stringent response mediated transcriptional reprogramming. A. Growth curve in M2G minimal media prior to 24 hours of starvation. Error bars represent +/- 1 SD of 3 biological replicates B. Growth curve in M2G minimal media following 24 hours of starvation. Error bars represent +/- 1 SD of 3 biological replicates C. Box-and-whisker plot of the ratio of CdnLDD to WT colony forming unit (CFU) ratio during competition. Dashed line indicates a ratio of 1, when number of CdnLDD colonies = number of WT colonies. Far left is nutrient-replete (M2G) conditions only; middle starts in nutrient-replete conditions; far right starts in carbon-starved (M2) conditions. Post-mix and pre-starve indicate time point just after mixing the strains and prior to the initial incubation. Center line in each box indicates mean of 6 biological replicates (3 mixed cultures in 2 combinations; for more detail, see Materials and Methods). Whiskers show minimum to maximum value. Box encases 25th to 75^th^ percentile.

Because we observed that cells with an inability to clear CdnL have a slight defect in outgrowth from carbon starvation, we wondered how this phenotype might relate to the effects of other transcriptional regulators involved in the stringent response. We repeated these experiments, this time assessing the growth of several mutant strains, including: a strain expressing an RNAP with a (p)ppGpp binding site 1 mutation (RNAP-1) (11); a strain lacking the transcription factor DksA (Δ*dksA*), which normally would bind to RNAP to form the second (p)ppGpp binding pocket during starvation (9, 12); a Δ*dksA*/ RNAP-1 strain, thus generating an RNAP that is essentially blind to the major effects of (p)ppGpp; and a CdnLDD/ Δ*dksA*/ RNAP-1 strain. Before starvation, the RNAP-1, Δ*dksA*, and Δ*dksA*/ RNAP-1 strains all exhibited growth rates very close to that of WT and CdnLDD, while the CdnLDD/ Δ*dksA*/ RNAP-1 strain had a slightly longer doubling time and lag time (Fig. 4A and Table S2). After the 24-hour starvation, there was a clear striation in outgrowth, with the RNAP-1 strain displaying a similar growth pattern to CdnLDD, followed by Δ*dksA* and Δ*dksA*/ RNAP-1. The CdnLDD/ Δ*dksA*/ RNAP-1 strain showed the greatest lag upon outgrowth, reaching an OD of 0.1 after about 11.6 hours in comparison to 8.3 hours for WT (Fig. 4B and Table S2). From these data, we conclude that the ability of *Caulobacter* to efficiently adapt to nutrient repletion following a period of starvation requires appropriate stringent response-mediated transcriptional reprogramming, which includes (p)ppGpp binding to RNAP, DksA, and clearance of CdnL.

### CdnL clearance during the SR is important for adaptation to nutrient limitation during competition

On its own, CdnL stabilization appeared to cause a slight defect in adaptation to changes in nutrient status (Fig. 4B and Table S2). We also observed transcriptional changes suggesting that CdnL stabilization promotes the transcription of genes that are typically downregulated upon SR activation (Fig. 3B – E). Because of these observations, we wondered if cells with a stabilized CdnL would be at a disadvantage if they were forced to compete with WT cells while adapting to changes in carbon availability. We tested this by competing a kanamycin-resistant or spectinomycin-resistant CdnLDD strain, which is unable to clear CdnL during starvation, with a spectinomycin-resistant or kanamycin-resistant WT strain, respectively. This allowed us to ask if the presence of CdnL is detrimental to fitness during starvation when in competition with cells that can effectively clear CdnL upon activation of the SR. We mixed equivalent OD units of the reciprocal strains in either M2G for nutrient-replete conditions or M2 for carbon-starved conditions. After 24 hours, we took a sample of the mixed culture and plated it on both a kanamycin plate and a spectinomycin plate in order to compare colony forming units (CFUs) between the CdnLDD and WT strains. We then either diluted the remaining culture into M2G to keep the cells in an exponential phase of growth, or into M2 to starve the cells.

We first tested the ability of CdnLDD to compete with WT under nutrient-rich conditions only. Over the course of 6 days, the CdnLDD and WT strains formed colonies in ∼1:1 ratio, suggesting that CdnL stabilization does not impact fitness when there are nutrients readily available (Fig. 4C, left). However, after periods of carbon starvation and recovery, the CdnLDD strains formed fewer colonies than the WT strains, indicating that the WT strains outcompeted the CdnLDD strains in the mixed culture (Fig. 4C, middle). This trend was observed regardless of the initial nutrient composition, and a competitive disadvantage was observed for CdnLDD after a single round of starvation (Fig. 4C, right). These data suggest that stabilization of CdnL puts cells at a disadvantage when trying to adapt to fluctuating nutrient availability and supports the idea that one way in which *Caulobacter* adapts to nutrient stress is through clearance of this transcriptional regulator.

## DISCUSSION

Adapting to environmental challenges such as nutrient deprivation requires efficient changes in transcription in order to downregulate biosynthetic genes and upregulate those that promote survival. While the direct binding of the SR alarmone (p)ppGpp to RNAP is a major way in which cells alter the transcriptome, other factors play key roles in allowing for an effective response to starvation. For instance, binding of the transcription factor DksA to RNAP has been shown to enhance the transcriptional effects of (p)ppGpp on RNAP (12); however, it is not only the presence of factors that promote stress responses that enables adaptation to nutrient stress, but also the absence of anabolic regulators that shifts the transcriptional balance from anabolism to survival.

Here, we show that *Caulobacter* CdnL, a CarD-family transcriptional regulator involved in ribosome biosynthesis and anabolism, is regulated upon SR activation (13–15). We find that transcriptional control at the *cdnL* promoter is not sufficient to control CdnL levels, and that regulation of CdnL protein occurs post-translationally in a manner dependent on SpoT, with (p)ppGpp being sufficient for CdnL clearance (23). We also show that CdnL is regulated at the protein level by the ClpXP protease, and that mutation of a degradation signal in the CdnL C-terminus stabilizes CdnL levels during the SR. Surprisingly, CdnLDD does not bind the chromosome after 60 minutes of starvation, nor does it globally alter transcription at this time point. Transcriptional changes that arise from CdnL stabilization are most obvious after 24 hours of starvation, where we find misregulation of ribosomal and metabolic genes. Ultimately, we find that clearance of CdnL is physiologically important, as CdnL stabilization causes adaptation defects. We propose that the combined and potentially synergistic actions of SpoT and ClpXP during the SR facilitate rapid clearance of CdnL protein. CdnL clearance allows for adaptation by transcriptionally downregulating ribosome biosynthesis and flux through metabolic pathways, thereby promoting *Caulobacter* survival when nutrients are lacking. When conditions become favorable again, re-introduction of CdnL enables cells to adapt by reactivating biosynthesis and metabolism, thereby allowing growth to resume.

The connection between the stringent response and the regulation of CdnL and its homologs in diverse bacteria has been a point of confusion. Indeed, transcription of mycobacterial CdnL homolog, *carD*, was initially shown to be upregulated during starvation, and CarD depletion was reported to sensitize cells to stressors such as starvation (20). However, recent studies indicate that while *carD* transcript levels increase due to stabilization of the transcript by the anti-sense RNA AscarD, CarD protein levels ultimately decrease through decreased translation and degradation by the Clp protease. Transcription of *ascarD* is proposed to be under control of a stress sigma factor SigF, thus connecting regulation of CarD to nutrient stress. This regulation of CarD was shown to help mycobacterial cells respond to various stresses (22).

Likewise, we find CdnL regulation to be functionally important for *Caulobacter* cells to adapt to the stress of nutrient fluctuations. Cells producing the stabilized CdnL variant, CdnLDD, have a slight defect upon outgrowth from carbon starvation, which is exacerbated in the absence of DksA and (p)ppGpp binding to RNAP (Fig. 4B). We also find that CdnLDD cells are outcompeted by WT cells when subjected to periods of carbon starvation, indicating that CdnL regulation is important for cells to adapt to nutrient stress (Fig. 4C).

These adaptation defects are likely caused by the effects of CdnLDD transcription (Fig. 3B - E). While we found that after 60 minutes of starvation, CdnLDD does not bind the chromosome and causes a relatively limited number of transcriptional changes, the effects of CdnL stabilization on transcription become more apparent after 24 hours of starvation and during the adaptation phase when nutrients are replenished. CdnL stabilization increases transcription of genes associated with ribosomes and metabolic pathways during nutrient stress. As ribosome biogenesis and maintenance are costly, it would be disadvantageous to promote these processes during times of limited resources. Likewise, promoting metabolic gene expression can lead to increased flux through these pathways during a time when extra metabolic activity would not be favored. Additionally, the redox reactions of the ETC, if inappropriately managed, can create damaging reactive oxygen species (ROS), which can cause DNA damage (33, 34). As many DNA damage response and repair proteins are upregulated in CdnLDD 60 minutes after the addition of glucose following a 24-hour starvation, it is tempting to speculate that increased flux through the ETC is creating ROS and causing DNA damage. The repair of damaged DNA coupled with the costly misregulation of ribosomes can put cells with a stabilized CdnL at a disadvantage when trying to adapt to nutrient fluctuations and reinitiate growth.

Regulation of CdnL during stress appears to be a common theme across diverse bacterial phyla. The *Borrelia burgdorferi* CdnL homolog, called LtpA, was found to be produced at 23⁰C, a condition that has been widely used to mimic *B. burgdorferi* in unfed ticks, while levels during incubation of cells at 37⁰C as well as during mammalian infection were significantly reduced (35). Deletion of *ltpA* prevented *B. burgdorferi* from infecting mice via tick infection, suggesting that LtpA could be important for *B. burgdorferi*’s survival within the tick vector and/or transmission to the mammalian host (35, 36). Similarly, depletion of the *Mycobacterium tuberculosis* homolog prevented cells from replicating and persisting in mice (20). *Bacillus cereus* homologs were found to be upregulated in response to various stressors and were important in the recovery-response to heat shock (37–39). Because CdnL homologs are broadly found and have been implicated in promoting adaptation and survival under different conditions, we believe that regulation of this transcription factor is a conserved mechanism enabling bacteria to adapt to stress.

## METHODS

### Caulobacter crescentus and Escherichia coli growth media and conditions

*C. crescentus* NA1000 cells were grown at 30°C in peptone-yeast extract (PYE) medium or minimal media described below. *E. coli* NEB Turbo (NEB Catalog #C2986K) and Rosetta(DE3)/pLysS cells were grown at 37°C and 30°C, respectively, in Luria-Bertani (LB) medium. Antibiotics for *Caulobacter* growth were used in liquid (solid) medium at the following concentrations: gentamycin, 1 (5) µg/mL; kanamycin, 5 (25) µg/mL; oxytetracycline, 1 (2) µg/mL; and spectinomycin, 25 (100) µg/mL. Streptomycin was used at 5 µg/mL in solid medium. *E. coli* antibiotics were used in liquid (solid) medium as follows: ampicillin, 50 (100) µg/mL; gentamicin, 15 (20) µg/mL; kanamycin, 30 (50) µg/mL; oxytetracycline, 12 (12) µg/mL; and spectinomycin, 50 (50) µg/mL. Strains and plasmids used in this study are listed in, Table S3.

### Starvation experiments

Cells were grown overnight in 2 – 4 mL minimal media with appropriate antibiotics. For both carbon and nitrogen starvation, M2G (M2 with 0.2% glucose w/v) was used (40). For phosphate starvation, Hutner base imidazole-buffered glucose glutamate medium (HIGG) was used (41). The next day, mid-log phase cells were harvested by centrifugation and washed thrice with nutrient-poor media. For carbon starvation, M2 media (without glucose) was used; for nitrogen starvation, M2G lacking NH_4_Cl was used; and for phosphate starvation, HIGG lacking Na_2_HPO_4_-KH_2_PO_4_ was used. After the final wash, cells were resuspended in 5 mL nutrient-limited media, divided into four 1.2 mL cultures, and incubated at 30⁰C shaking at 225 rpm. OD_600_ was taken every 30 minutes from 0 to 90 minutes. 0.5 or 1 mL cells were harvested by centrifugation and resuspended in OD_600_/0.003 or OD_600_/0.003 uL 1X SDS loading dye, respectively, for subsequent immunoblotting. Three biological replicates were obtained for each strain in each condition.

### RelA’ induction

EG1799 and EG1800 were grown overnight in 5 mL M2G with appropriate antibiotics. The next day, mid-log phase cells were diluted to OD_600_ = 0.2 in 10 mL M2G with 0.3% xylose (w/v). Cultures were distributed into seven 1.4 mL aliquots and incubated as described above. OD_600_ was taken every 30 minutes from 0 to 120 minutes and protein samples for immunoblotting were made by resuspension in 1X SDS loading dye. Three biological replicates were obtained for each strain.

### Transcriptional shut-off

EG3190, EG3193, and EG3194 were grown overnight in 2 – 4 mL M2 with 0.3% xylose and appropriate antibiotics. The next day, mid-log phase cells were harvested by centrifugation and washed thrice with M2G (to shut off transcription) or M2 (to shut off transcription and starve cells). After the final wash, cells were resuspended in 5 mL M2G or M2 and were incubated as described above. A total of 3 biological replicates were obtained for each strain. Immunoblotting samples were prepared by resuspension in 1X SDS loading dye.

### ClpX* induction

EG802 and EG865 were grown overnight in 2 – 4 mL M2G with appropriate antibiotics. The next day, mid-log phase cultures were diluted to OD_600_ = 0.1 – 0.25 in 5 mL M2G supplemented with 0.3% xylose (w/v) and grown for 2 hours. The starvation was performed as described above. Three biological replicates were obtained for each strain.

### Immunoblotting

Equivalent OD units of cell lysate were loaded on an SDS-PAGE gel following cell harvest by centrifugation, lysing 1X SDS loading dye, and boiling for 5 – 10 minutes. SDS-PAGE and protein transfer to nitrocellulose membranes were followed using standard procedures. Antibodies were used at the following dilutions: CdnL 1:10,000 (14); MreB 1:10,000 (Régis Hallez, University of Namur); SpmX 1:20,000 (Patrick Viollier, University of Geneva); GFP 1:2,000 (Clonetech Labs Catalog #NC9777966); HRP-labeled α-rabbit secondary 1:10,000 (BioRAD Catalog #170-6515); and/or HRP-labeled α-mouse secondary (Cell Signaling Technology Catalog #7076S). Clarity Western Electrochemiluminescent substrate (BioRAD Catalog #170-5060) was used to visualize proteins on an Amersham Imager 600 RGB gel and membrane imager (GE).

### Protein purification

CdnL and CdnLDD were overproduced in Rosetta (DE3) pLysS *E. coli* from pEG1129 (His-SUMO-CdnL) and pEG1634 (His-SUMO-CdnLDD), respectively. Cells were induced with 1mM IPTG for 3 hours at 30°C. Cell pellets were resuspended in Column Buffer A (50 mM Tris-HCl pH 8.0, 300 mM NaCl, 10% glycerol, 20 mM imidazole, 1 mM β-mercaptoethanol), flash frozen in liquid nitrogen, and stored at −80°C. To purify the His-SUMO tagged proteins, pellets were thawed at 37°C, and 10 U/mL DNase I, 1 mg/mL lysozyme, and 2.5 mM MgCl2 were added. Cell slurries were rotated at room temperature for 30 minutes, then sonicated and centrifuged for 30 minutes at 15,000 x g at 4°C. Protein supernatants were then filtered and loaded onto a pre-equilibrated HisTrap FF 1mL column (Cytiva, Marlborough, Massachusetts). His-SUMO-CdnL and His-SUMO-CdnLDD were eluted in 30% Column Buffer B (same as Column Buffer A but with 1M imidazole), and peak fractions were concentrated. The His-Ulp1 SUMO protease was added to a molar ratio (protease:protein) of 1:290 and 1:50 for His-SUMO-CdnL and His-Sumo-CdnLDD, respectively, and dialyzed into 1 L Column Buffer A overnight at 4°C. The cleaved protein solutions were again loaded onto a HisTrap FF 1mL column. Peak flowthrough fractions were combined, concentrated, and applied to a Superdex 200 10/300 GL (Cytiva) column equilibrated with storage buffer (50 mM HEPES-NaOH pH 7.2, 100 mM NaCl, 10% glycerol). Peak fractions were combined, concentrated, snap-frozen in liquid nitrogen, and stored at −80°C. ClpX was purified using a similar protocol as described for *E. coli* ClpX (24). Recombinant his-tagged ClpP was purified as described (42).

### *In vitro* proteolysis assays

*In vitro* proteolysis reactions were performed with purified ClpX, ClpP, and CdnL or CdnLDD as previously described (25).

### Monitoring CdnL protein levels

EG865, EG1139, and EG2530 were inoculated in 2 mL M2G. The next day, the cultures were diluted to an OD_600_ of ∼0.001 and grown for approximately 18 hours or until the cultures reached an OD_600_ of ∼0.4 – 0.5. At OD_600_ of ∼0.4 – 0.5, 500 μL of culture sample was taken to make protein samples for immunoblotting as described above. Samples were then taken in the following 1, 2, 4, 6, 7, 8, and 25 hours. Immunoblotting was performed as described above, except following the transfer, membranes were stained with Ponceau stain for total protein normalization.

### Chromatin immunoprecipitation coupled to deep sequencing (ChIP-seq)

EG865, EG1898, and EG2530 were grown overnight in 6 mL M2G (M2 with 0.2% glucose w/v) at 30°C (shaking at 200 rpm), respectively. The next day, cultures were diluted into 100 mL M2G and grown to an OD_660_ of 0.5. Cells were harvested by centrifugation at 8,000 rpm for 10 min at 25°C, and cell pellets were washed 3 times with 50 mL of M2 media (without glucose) at 25°C. Each of the washed cell pellets was split into two and used to inoculate 50 mL of M2 (carbon starvation) and 50 mL of M2G (preheated culture medium at 30°C), respectively. Cultures were incubated at 30⁰C (shaking at 200 rpm) for additional 60 minutes and then, supplemented with 10 μM sodium phosphate buffer (pH 7.6) and treated with formaldehyde (1% final concentration) at RT for 10 min to achieve crosslinking. Subsequently, the cultures were incubated for an additional 30 min on ice and washed three times in phosphate buffered saline (PBS, pH 7.4). The resulting cell pellets were stored at −80°C. After resuspension of the cells in TES buffer (10 mM Tris-HCl pH 7.5, 1 mM EDTA, 100 mM NaCl) containing 10 mM of DTT, the cell resuspensions were incubated in the presence of Ready-Lyse lysozyme solution (Epicentre, Madison, WI) for 10 minutes at 37°C, according to the manufacturer’s instructions. Lysates were sonicated (Bioruptor Pico) at 4°C using 15 bursts of 30 sec to shear DNA fragments to an average length of 0.3–0.5 kbp and cleared by centrifugation at 14,000 rpm for 2 min at 4°C. The volume of the lysates was then adjusted (relative to the protein concentration) to 1 mL using ChIP buffer (0.01% SDS, 1.1% Triton X-84 100, 1.2 mM EDTA, 16.7 mM Tris-HCl [pH 8.1], 167 mM NaCl) containing protease inhibitors (Roche) and pre-cleared with 80 μL of Protein-A agarose (Roche, www.roche.com) and 100 μg BSA. Five percent of the pre-cleared lysates were kept as total input samples. The rest of the pre-cleared lysates were then incubated overnight at 4°C with polyclonal rabbit antibodies targeting CdnL (1:1,000 dilution). The Immuno-complexes were captured by incubation with Protein-A agarose beads (pre-saturated with BSA) during a 2 hr incubation at 4°C and then, washed subsequently with low salt washing buffer (0.1% SDS, 1% Triton X-100, 2 mM EDTA, 20 mM Tris-HCl pH 8.1, 150 mM NaCl), with high salt washing buffer (0.1% SDS, 1% Triton X-100, 2 mM EDTA, 20 mM Tris-HCl pH 8.1, 500 mM NaCl), with LiCl washing buffer (0.25 M LiCl, 1% NP-40, 1% deoxycholate, 1 mM EDTA, 10 mM Tris-HCl pH 8.1) and finally twice with TE buffer (10 mM Tris-HCl pH 8.1, 1 mM EDTA). The immuno-complexes were eluted from the Protein-A beads with two times 250 μL elution buffer (SDS 1%, 0.1 M NaHCO3, freshly prepared) and then, just like the total input sample, incubated overnight with 300 mM NaCl at 65°C to reverse the crosslinks. The samples were then treated with 2 μg of Proteinase K for 2 hr at 45°C in 40 mM EDTA and 40 mM Tris-HCl (pH 6.5). DNA was extracted using phenol:chloroform:isoamyl alcohol (25:24:1), ethanol-precipitated using 20 μg of glycogen as a carrier and resuspended in 30 μl of DNAse/RNAse free water.

Immunoprecipitated chromatins were used to prepare sample libraries used for deep-sequencing at Fasteris SA (Geneva, Switzerland). ChIP-Seq libraries were prepared using the DNA Sample Prep Kit (Illumina) following manufacturer instructions. Single-end run was performed on an Illumina Next-Generation DNA sequencing instruments (NextSeq High), 50 cycles were performed and yielded several million reads per sequenced samples. The single-end sequence reads stored in FastQ files were mapped against the genome of *C. crescentus* NA1000 (NC_011916.1) using Bowtie2 Version 2.4.5+galaxy1 available on the web-based analysis platform Galaxy (https://usegalaxy.org) to generate the standard genomic position format files (BAM).

ChIP-Seq reads sequencing and alignment statistics are summarized in DatasetS1. Then, BAM files were imported into SeqMonk version 1.47.2 (http://www.bioinformatics.babraham.ac.uk/projects/seqmonk/) to build ChIP-Seq normalized sequence read profiles. Briefly, the genome was subdivided into 50 bp, and for every probe, we calculated the number of reads per probe as a function of the total number of reads (per million, using the Read Count Quantitation option; DatasetS1).

Using the web-based analysis platform Galaxy (https://usegalaxy.org), CdnL(M2+G) ChIP-Seq peaks were called using MACS2 Version 2.2.7.1+galaxy0 (No broad regions option) relative to the *ΔcdnL*(M2+G) negative ChIP-seq control. The q-value (false discovery rate, FDR) cut-off for called peaks was 0.05. Peaks were rank-ordered according to their fold-enrichment values (DatasetS1, the 107 ChIP-Seq CdnL(M2+G) statistical peaks with a fold enrichment greater than 2 were retained for further analysis). MACS2 analysed data illustrated in Figure S5 (CdnL relevant peaks identified by ChIP-seq) are provided in DatasetS1. Then, CdnL(M2+G) ChIP-seq equivalent peaks (overlapping peak “start” and “end” MACS2 coordinates) were searched in the MACS2 statistical peak analyses of ChIP-Seq CdnL(M2-G), ChIP-Seq CdnLDD(M2+G), and ChIP-Seq CdnLDD(M2-G) datasets (relative to the *ΔcdnL*(M2+G) or (M2-G) negatives ChIP-seq controls, accordingly). In cases where no overlapping peaks were identified, a default fold enrichment value of 1 was assigned. To facilitate visualization and ChIP-seq analysis, these data were subsequently submitted to the http://www.heatmapper.ca/expression/ website to generate a heat map (Parameters: Clustering method “average linkage”; Distance measurement method “Pearson”; Scale “Row Z-score”). Row Z-score = (CdnL Fold enrichment value in the sample of interest - Mean CdnL Fold enrichment across all samples) / Standard Deviation. Analysed data illustrated in Figure 3A as well as the list of genes composing each group are provided in DatasetS1. For each group defined during the heat map analysis, the list of genes potentially regulated by CdnL (presence of a CdnL statistically significant peak detected on the gene’s promoter region) was submitted to the DAVID website (The Database for Annotation, Visualization and Integrated Discovery; https://david.ncifcrf.gov/home.jsp) to identify enriched biological themes, particularly GO terms (DatasetS1). Sequence data have been deposited to the Gene Expression Omnibus (GEO) database (GSE249185 series, accession numbers GSM7927858 – GSM7927866).

### RNA sequencing (RNA-seq)

For 0 and 60 minute starvation samples, cultures of 3 independent colonies of EG865 and EG2530 were inoculated into 2 mL M2G overnight. The next day, cultures were diluted into 4 mL M2G and grown to an OD_600_ of 0.4 – 0.6. Cells were prepared for carbon starvation as described above, except after the final wash, cells were resuspended in 4 mL M2 and were split into two 2 mL samples each. One sample for each replicate was incubated at 30⁰C shaking at 225 rpm for 60 minutes, while the other was stabilized using RNAprotect Bacteria Reagent (Qiagen Catalog #76506) following manufacturer’s instructions. Briefly, 4 mL RNAprotect was added to 15 mL conical tubes, to which the 2 mL culture samples were added. The conical tubes were vortexed for 5 seconds, incubated at room temperature for 5 minutes, and then centrifuged for 10 minutes at 5,000 x g. The pellets were flash frozen in liquid nitrogen and stored at −80°C. The 60-minute starved samples were harvested in the same way.

For 24 hour starved and 60 minute recovery samples, cultures of 3 independent colonies were inoculated into 6 mL M2G and grown overnight. Once cultures were at an OD_600_ of 0.4 – 0.6, cultures were prepared for carbon starvation as described above except after final wash, cells were resuspended in 6 mL M2 and incubated at 30⁰C shaking at 225 rpm for 24 hours. After 24 hours, 2 mL of each culture was removed and stabilized using RNAprotect as described above and flash frozen in liquid nitrogen. To the remaining 4 mL of culture, 40 μL of 20% glucose solution was added, and cultures were incubated for 60 minutes. Recovery samples were stabilized with RNAprotect and flash frozen in liquid nitrogen.

All samples were processed at SeqCenter (Pittsburg,PA). There, samples were DNAse treated with Invitrogen DNAse (RNAse free). Library preparation was performed using Illumina’s Stranded Total RNA Prep Ligation with Ribo-Zero Plus kit and 10bp IDT for Illumina indices and *Caulobacter*-specific rRNA depletion probes. Sequencing was done on a NextSeq2000 giving 2×51bp reads. Quality control and adapter trimming was performed with bcl2fastq (version 2.20.0.445; default parameters). Read mapping was performed with HISAT2 (version 2.2.0; default parameters + ‘--very-sensitive’) (43). Read quantification was performed using Subread’s featureCounts functionality (version 2.0.1; default parameters + ‘-Q 20’) (44). Read counts were loaded into R (version 4.0.2; default parameters) and were normalized using edgeR’s (version 1.14.5; default parameters) Trimmed Mean of M values (TMM) algorithm (45). Subsequent values were then converted to counts per million (cpm). Differential expression analysis was performed using edgeR’s exact test for differences between two groups of negative-binomial counts. All RNA-seq data have been deposited in the Sequence Read Archive (SRA) under accession numbers SRR27130076 - SRR27130078, SRR27130555 - SRR27130557, and SRR27146334 - SRR27146351, and are associated with BioProject PRJNA1049818.

### Starvation outgrowth growth curves

EG865, EG2530, EG2652, EG2756, EG2776, and EG2861 were grown overnight in 2 mL M2G. The next day, biological triplicates were diluted to OD_600_ = 0.05 in 100 µL M2G in a 96 well plate. A Cytation1 imaging reader (Agilent, Biotek) measured absorbance every 30 minutes for 36 hours with intermittent shaking. The remainder of each 2 mL culture was back-diluted into 5 mL M2G and incubated overnight. The next day, log-phase cells were harvested by centrifugation at 8,250 x g for 5 minutes, washed thrice with M2, resuspended in a total volume of 4.5 mL M2, and incubated for 24 hours. Following the 24-hour starvation, another growth curve in M2G was run as described above. Approximate time to OD = 0.1 was calculated by solving a trendline using averaged triplicates just prior to OD 0.1 and just after OD 0.1. Doubling times were calculated using GraphPad Prism software.

### Competition experiments

EG3402, EG3404, EG3406, and EG3408 were grown overnight in 2 mL PYE with appropriate antibiotics. The next day, overnight cultures were pelleted at 16,300 x g for one minute and washed twice with M2G. Following the final wash, cells were resuspended in 10 mL M2G with appropriate antibiotics and grown overnight. The next day, stationary phase cells were diluted into 50 mL M2G with appropriate antibiotics and allowed to grow a minimum of 6 hours (approximately 2 doublings in M2G) to an OD_600_ of 0.3 – 0.6. Cells were then synchronized with modifications to the original protocol (46). Briefly, cells were pelleted at 6,000 x g for 10 minutes and then resuspended in 1.5 mL cold 1X M2 salts. The resuspension was transferred to 15 mL Corex tubes. 1.5 mL cold Percoll was added and cells were centrifuged at 15,000 x g for 20 minutes at 4°C. Swarmer cells were transferred to 15 mL conicals, 1X M2 salts were added to fill the tube, and cells were pelleted at 10,000 x g for 5 minutes. Cell pellets were resuspended in 1 mL 1X M2 salts and centrifuged at 16,300 x g for 1 minute. The resulting pellet was resuspended in M2 for nutrient-poor conditions or M2G for nutrient-rich conditions and OD_600_ was recorded.

When starting in nutrient-poor (M2) conditions, cultures were equally diluted to OD_600_ = 0.25 - 0.4 and mixed 1:1 (EG3402: EG3408, EG3404: EG3406) in 5 mL M2 and incubated for 24 hours. Prior to incubation, 50 μL of the mixed cultures were serially diluted, and 100 μL of 10^-4^ – 10^-6^ dilutions were plated on both PYE-spectinomycin and PYE-kanamycin plates. When starting in nutrient-rich conditions, the same procedure was followed except after plating, the mixed cultures were diluted to OD ∼ 0.0001 and 0.00005 in 5 mL M2G and allowed to grow for 24 hours.

Every 24 hours, a culture sample was taken for plating as described above. Colony forming units (CFUs) were counted on plates following a 2-day incubation at 30⁰C, and a ratio called the competitive index was calculated from the CFUs formed by the CdnLDD strains compared to the WT strains (EG3408/EG3402; EG3406/EG3404) on the plates that yielded the maximum number of countable colonies.

If in nutrient-rich conditions, a volume of cells approximately equal to OD_600_ = 0.25 were pelleted, washed twice with M2, resuspended in 5 mL M2, and incubated for a 24-hour starvation. If in nutrient-poor conditions, cells were diluted to an OD ∼ 0.0003 in 5 mL M2G and incubated for a 24-hour recovery. This was repeated for a total of three rounds. The entire process was repeated for a total of 4 biological replicates for the competition starting in nutrient-poor conditions and 3 biological replicates for the competition starting in nutrient-rich conditions. For the competition in only nutrient-rich conditions, a similar procedure was followed, but every 24 hours cells were diluted to an OD ∼ 0.00005 in 5 mL M2G for a total of 6 rounds. The entire process was repeated to obtain 3 biological replicates.

## ACKNOWLEDGEMENTS

We thank Allison Daitch and Marie Stoltzfus for the construction of some *Caulobacter* strains and plasmids. We thank Allen Buskirk and Annie Campbell for helpful discussions about ribosomes. This project was supported by funds from the NIH (NIH-NIGMS Grant R35GM136221 to E.D.G. and Grant R35GM130320 to P.C.) and the Swiss National Science Foundation (SNSF) (project grant 310030_212531 to P.H.V). E.S. was supported in part through the Biochemistry, Cellular, and Molecular Biology training program (NIH-NIGMS Grant T32GM144272).

---

**Supplemental Figure S1:**
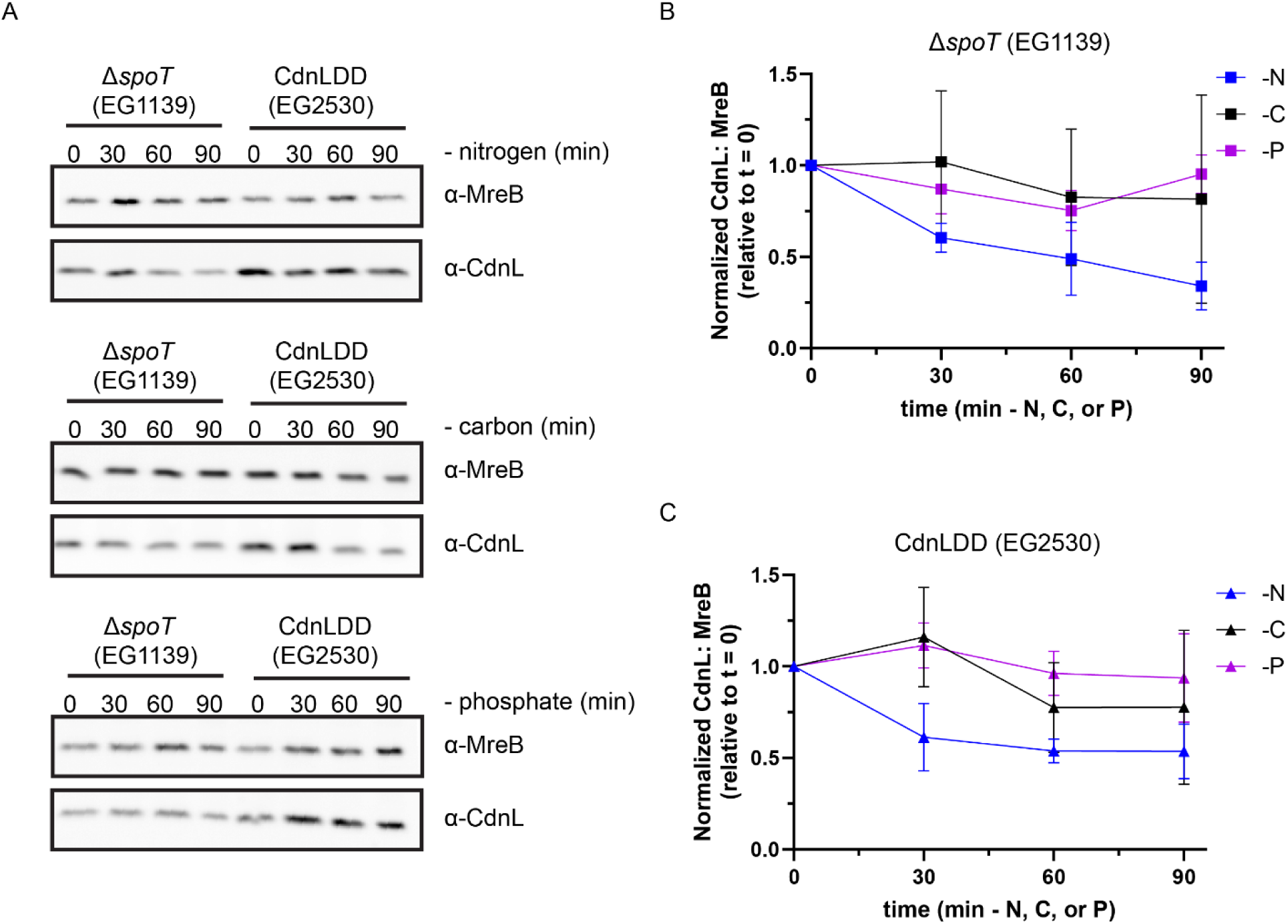
CdnL levels are stabilized in ΔspoT and CdnLDD. A. Representative western blot of CdnL during 90 minutes of nitrogen, carbon, and phosphate starvation in Δ*spoT* (EG1139) and CdnLDD (EG2530). Protein samples were taken every 30 minutes. MreB was used as a loading control. B. Densitometry of CdnL levels (normalized to MreB) relative to t = 0 during nitrogen (-N), carbon (-C), and phosphate (-P) starvations in Δ*spoT* (EG1139) from western blots as performed in A. Error bars represent +/- 1 SD of 3 biological replicates. C. Densitometry of CdnL levels (normalized to MreB) relative to t = 0 during nitrogen (-N), carbon (-C), and phosphate (-P) starvations in CdnLDD (EG2530) from western blots as performed in A. Error bars represent +/- 1 SD of 3 biological replicates.

**Supplemental Figure S2:**
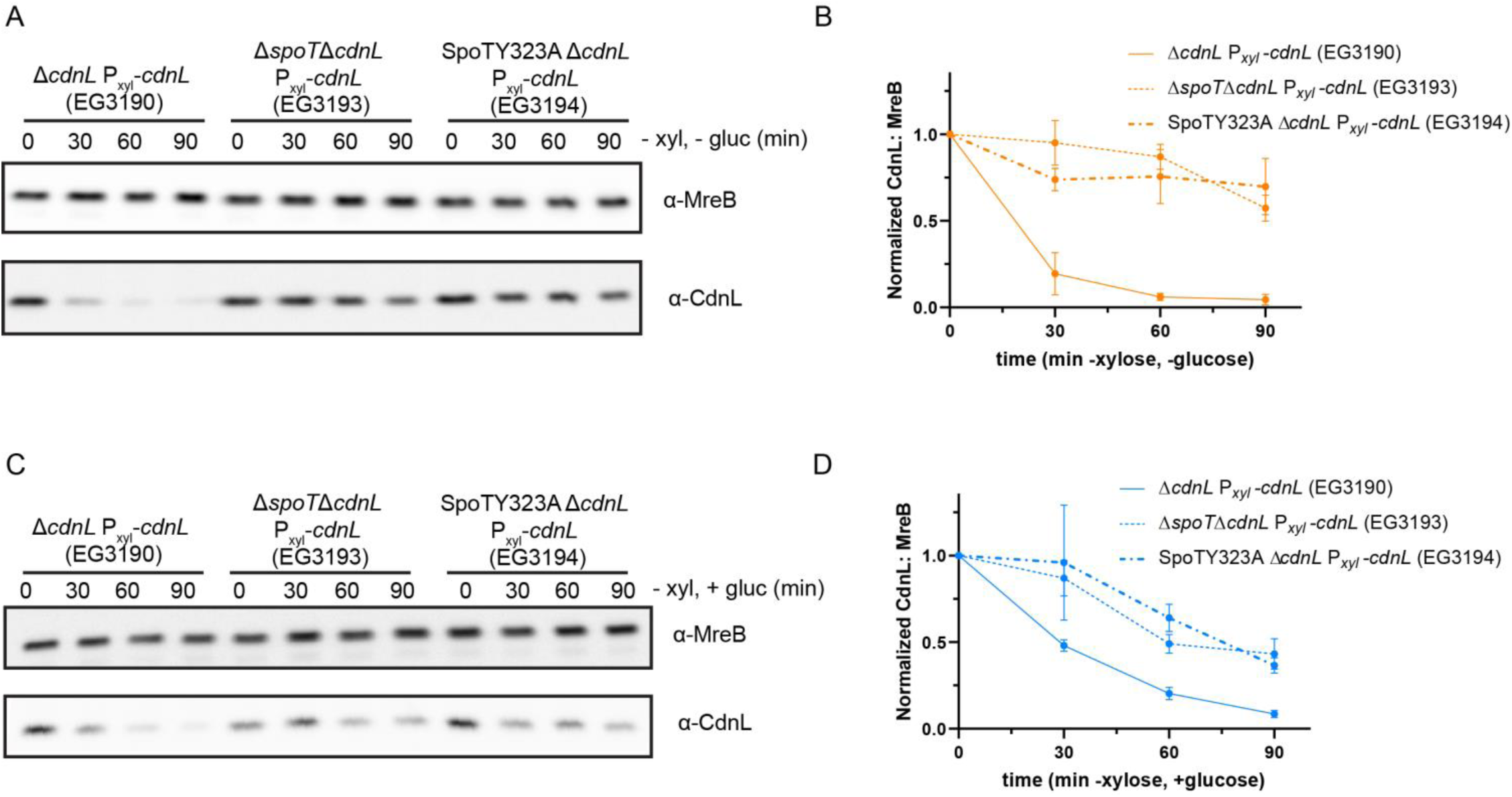
Transcriptional regulation of *cdnL* is not sufficient to control CdnL levels. A. Representative western blot of CdnL during 90 minutes of glucose (gluc) starvation with xylose (xyl) depletion of *cdnL* in a Δ*cdnL,* Δ*spoT*Δ*cdnL*, or SpoTY323AΔ*cdnL* background. Protein samples were taken every 30 minutes. MreB was used as a loading control. B. Densitometry of CdnL levels (normalized to MreB) relative to t = 0 from western blots as performed in A. Error bars represent +/- 1 SD of 3 biological replicates. C. Representative western blot of CdnL during xylose (xyl) depletion of *cdnL* in a Δ*cdnL,* Δ*spoT*Δ*cdnL*, or SpoTY323AΔ*cdnL* background. Protein samples were taken every 30 minutes. MreB was used as a loading control. D. Densitometry of CdnL levels (normalized to MreB) relative to t = 0 from western blots as performed in C. Error bars represent +/- 1 SD of 3 biological replicates.

**Supplemental Figure S3:**
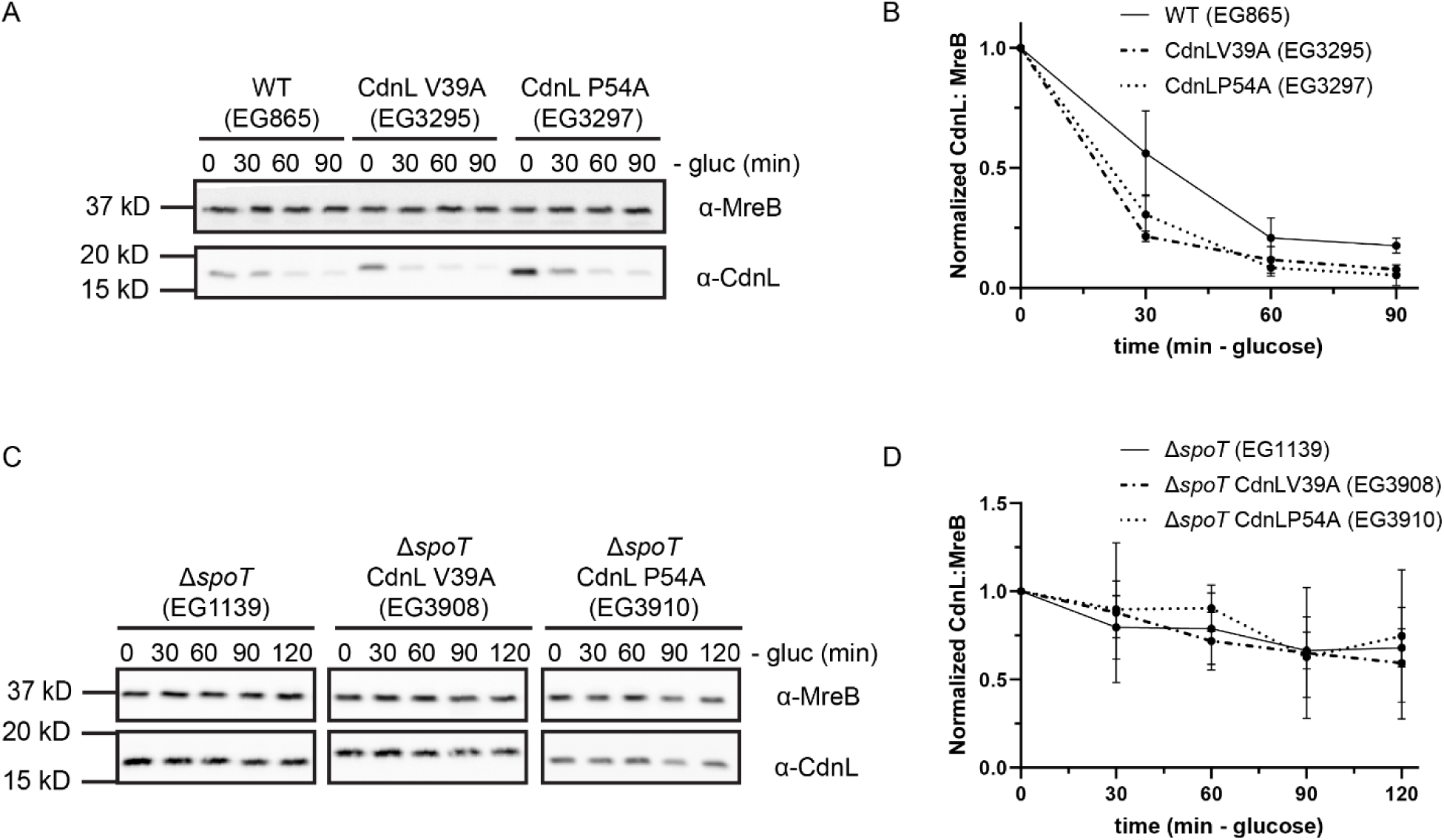
CdnL mutants with a reduced interaction with RNAP are cleared more rapidly than WT CdnL. A. Representative western blot of WT CdnL, CdnL V39A, or CdnL P54A during 90 minutes of glucose (gluc) starvation. Protein samples were taken every 30 minutes. MreB was used as a loading control. B. Densitometry of WT CdnL, CdnL V39A, or CdnL P54A levels (normalized to MreB) relative to t = 0 from western blots as performed in A. Error bars represent +/- 1 SD of 3 biological replicates. C. Representative western blot of WT CdnL, CdnL V39A, or CdnL P54A in a Δ*spoT* background during 90 minutes of glucose (gluc) starvation. Protein samples were taken every 30 minutes. MreB was used as a loading control. D. Densitometry of GFP-AA levels (normalized to SpmX) relative to t = 0 from western blots as performed in A. Error bars represent +/- 1 SD of 3 biological replicates.

**Supplemental Figure S4:**
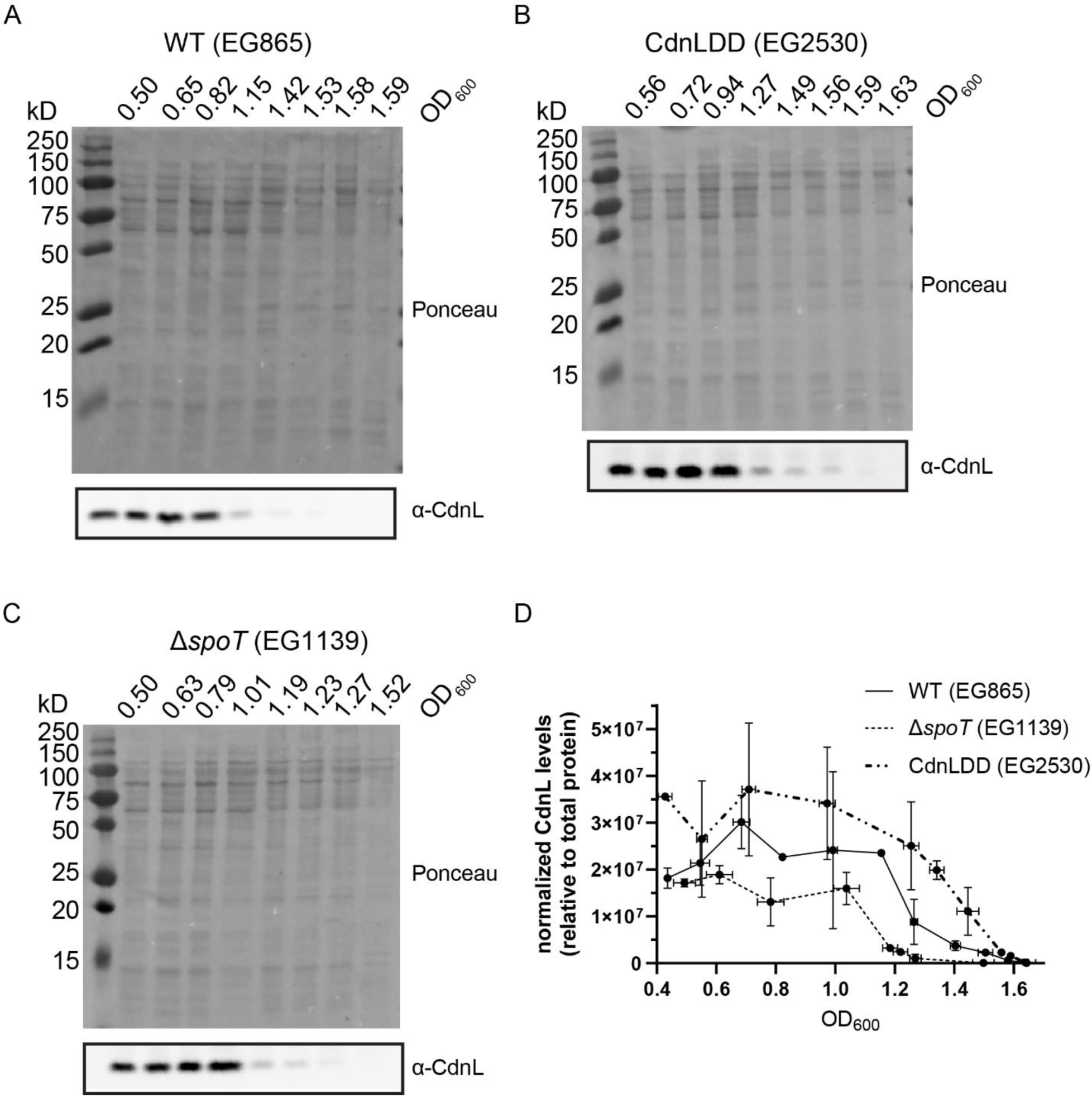
CdnL clearance during stationary phase is SpoT- and ClpXP-independent. A. Representative Ponceau stain and western blot for CdnL in WT (EG865). Samples were taken once OD_600_ = 0.4 – 0.6 for t = 0, and then at t = 1, 2, 4, 6, 7, 8, and 25 hours later. OD_600_ was recorded at each timepoint. B. Representative Ponceau stain and western blot for CdnL in Δ*spoT* (EG1139). Samples were taken once OD_600_ = 0.4 – 0.6 for t = 0, and then at t = 1, 2, 4, 6, 7, 8, and 25 hours later. OD_600_ was recorded at each timepoint. C. Representative Ponceau stain and western blot for CdnL in CdnLDD (EG2530). Samples were taken once OD_600_ = 0.4 – 0.6 for t = 0, and then at t = 1, 2, 4, 6, 7, 8, and 25 hours later. OD_600_ was recorded at each timepoint. D. Densitometry analysis of CdnL/CdnLDD (normalized to total protein) for WT, Δ*spoT,* or CdnLDD strains from OD_600_ ∼ 0.4 – 1.6 using Ponceau-stained membranes and western blots as performed in A - C. CdnL levels for OD_600_ values within 0.1 units were averaged. X and Y error bars represent +/- 1 SD of 1 – 3 biological replicates per point.

**Supplemental Figure S5:**
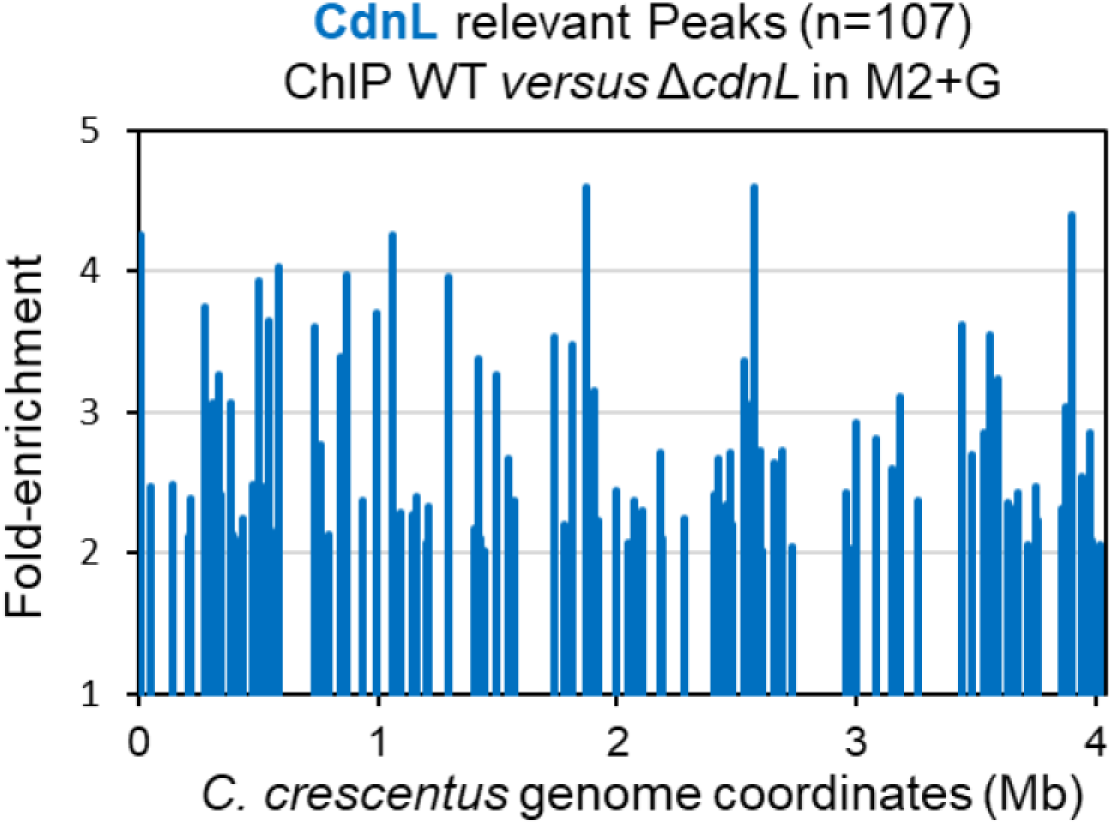
CdnL relevant peaks identified by ChIP-seq. 107 peaks identified by ChIP-seq as having a > 2-fold enrichment compared to the Δ*cdnL* (EG1898) control in M2G. These peaks were selected for further analyses in comparing WT (EG865) to CdnLDD (EG2530) in M2G and M2.

**Supplemental Figure S6:**
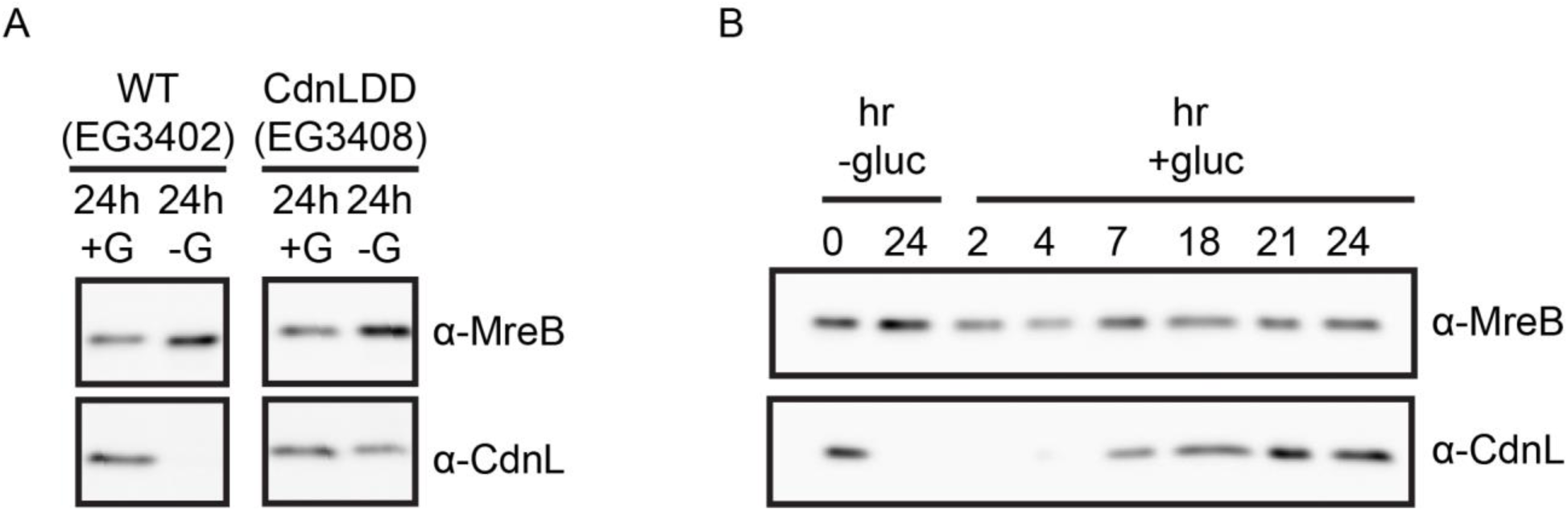
CdnLDD is stable after 24 hours of starvation. A. Representative western blot of WT (EG3402) and CdnLDD (EG3408) before (24h +G) and after (24h -G) 24 hours of starvation. MreB was used as a loading control. B. Representative western blot of CdnL levels in WT (EG3402) after 24 hours of starvation and at indicated timepoints following the addition of glucose. MreB was used as a loading control.

