## Supplementary material for "Regulation of the transcription factor CdnL promotes adaptation to nutrient stress in *Caulobacter*": SI Appendix

**Supplemental Table S1: Half-life values**

| **Strain/condition** | **Half-life (min)** |
|---|---|
| WT (EG865) -C | 15 ± 2 |
| Δ*spoT* (EG1139) -C | > 500 |
| CdnLDD (EG2530) -C | > 500 |
| SpoTY323A (EG2221) -C | > 500 |
| WT (EG865) -N | 18 ± 1 |
| Δ*spoT* (EG1139) -N | 60 ± 20 |
| CdnLDD (EG2530) -N | 87 ± 19 |
| WT (EG865) -P | > 500 |
| Δ*spoT* (EG1139) -P | > 500 |
| CdnLDD (EG2530) -P | > 500 |
| RelA' (EG1799) | 103 ± 16 |
| RelA'-dead (EG1800) | > 500 |
| Δ*cdnL* $P_{xyl}$-*cdnL* (EG3190) -xyl, -gluc | 13 ± 4 |
| Δ*spoT*Δ*cdnL* $P_{xyl}$-*cdnL* (EG3193) -xyl, -gluc | 139 ± 21 |
| SpoTY323A Δ*cdnL* $P_{xyl}$-*cdnL* (EG3194) -xyl, -gluc | 334 ± 288 |
| Δ*cdnL* $P_{xyl}$-*cdnL* (EG3190) -xyl | 27 ± 2 |
| Δ*spoT*Δ*cdnL* $P_{xyl}$-*cdnL* (EG3193) -xyl | 70 ± 12 |
| SpoTY323A Δ*cdnL* $P_{xyl}$-*cdnL* (EG3194) -xyl | 75 ± 8 |
| $P_{ruv}$-GFP-AA (EG3898) | 14 ± 3 |
| Δ*spoT* $P_{ruv}$-GFP-AA (EG3899) | 68 ± 25 |
| WT (EG865) | 32 ± 7 |
| CdnLV39A (EG3295) | 15 ± 1 |
| CdnLP54A (EG3297) | 18 ± 3 |

Error indicates +/- 1 SD of 3 biological replicates

**Supplemental Table S2: Outgrowth and doubling times for Figure 3**

| | Strain | Approx. time to OD = 0.1 (hr) | $t_d$ (hr) |
|---|---|---|---|
| Pre-starve | WT (EG865) | 4.2 | 2.1 ± 0.06 |
| | CdnLDD (EG2530) | 4.5 | 2.2 ± 0.02 |
| | RNAP-1 (EG2652) | 4.5 | 2.2 ± 0.02 |
| | Δ*dksA* (EG2756) | 4.0 | 2.1 ± 0.15 |
| | RNAP-1/Δ*dksA* (EG2776) | 4.8 | 2.3 ± 0.02 |
| | CdnLDD/RNAP-1/Δ*dksA* (EG2861) | 5.5 | 2.5 ± 0.24 |
| Post-starve | WT (EG865) | 8.3 | 1.9 ± 0.14 |
| | CdnLDD (EG2530) | 9.3 | 2.1 ± 0.02 |
| | RNAP-1 (EG2652) | 9.1 | 2.2 ± 0.09 |
| | Δ*dksA* (EG2756) | 10.3 | 2.6 ± 0.19 |
| | RNAP-1/Δ*dksA* (EG2776) | 10.5 | 2.9 ± 0.52 |
| | CdnLDD/RNAP-1/Δ*dksA* (EG2861) | 11.6 | 2.7 ± 0.05 |

Error indicates +/- 1 SD of 3 biological replicates

**Supplemental Table S3: Plasmids and Strains used in this study**

**Dataset S1: ChIP-seq data of WT CdnL and CdnLDD in M2G and 60 minutes in M2**

**Dataset S2: RNA-seq comparing WT (EG865) to CdnLDD (EG2530) at 0 minutes of starvation**
Includes all quantified genes, genes > 2-fold differentially regulated ($p < 0.05$ and FDR < 0.05), and DAVID analyses

**Dataset S3: RNA-seq comparing WT (EG865) to CdnLDD (EG2530) at 60 minutes of starvation**
Includes all quantified genes, genes > 2-fold differentially regulated ($p < 0.05$ and FDR < 0.05), and DAVID analyses

**Dataset S4: RNA-seq data comparing WT (EG865) 60 minutes starved to 24 hours starved**
Includes all quantified genes and genes > 2-fold differentially regulated ($p < 0.05$ and FDR < 0.05)

**Dataset S5: RNA-seq comparing WT (EG865) to CdnLDD (EG2530) at 24 hours of starvation**

Includes all quantified genes, genes > 2-fold differentially regulated ($p < 0.05$ and FDR < 0.05), and DAVID analyses

**Dataset S6: RNA-seq comparing WT (EG865) to CdnLDD (EG2530) at 60 minutes of recovery after glucose addition**

Includes all quantified genes, genes > 2-fold differentially regulated ($p < 0.05$ and FDR < 0.05), and DAVID analyses
